# Systematic multivariate analysis of chromatin complex dependencies reveals Set1C/COMPASS as a melanoma-enriched epigenetic vulnerability

**DOI:** 10.64898/2026.02.13.705694

**Authors:** Luisa Quesada Camacho, Mohammad Fallahi-Sichani

## Abstract

Epigenetic dysregulation is a common feature of cancer. It creates vulnerabilities arising from an increased reliance on chromatin-based mechanisms that sustain malignant transcriptional states. While many chromatin regulators are broadly required for cellular viability, others function in a context-dependent manner across distinct oncogenic settings, tissue lineages, and differentiation states. Moreover, chromatin regulators often operate within multi-subunit complexes; thus, epigenetic vulnerabilities emerge from coordinated complex activities. Here, we integrate large-scale genetic dependency maps from human cancer cell lines with curated epigenetic complex annotations to perform a systematic, multivariate analysis of complex-level epigenetic dependencies across cancer lineages. Our analysis reveals that dependencies frequently cluster among functionally related chromatin complexes and that biologically related cancer types (e.g., hematologic malignancies) share similar dependency patterns, consistent with shared underlying epigenetic requirements. Focusing on melanoma, we identify multiple enriched epigenetic complex dependencies, including complexes previously associated with recurrent genetic alterations or melanocyte lineage regulation, as well as a previously unrecognized vulnerability involving the H3K4 methyltransferase complex Set1C/COMPASS. This dependency is not restricted to a specific melanoma differentiation state, but genetic depletion of CXXC1 (a critical complex-specific subunit) suggests that CXXC1-dependent melanoma cells require Set1C/COMPASS activity to maintain global H3K4 trimethylation (H3K4me3) and proliferation. Integrative modeling links Set1C/COMPASS dependency to MYC- and E2F-driven transcriptional programs, which are suppressed upon complex inhibition. Together, this work combines integrative, multivariate analysis of lineage-enriched epigenetic dependencies with genetic perturbation, transcriptional profiling, and single-cell analysis to uncover an enriched epigenetic dependency on Set1C/COMPASS in melanoma cells.

**Author Summary:** Cancer cells often rely on abnormal regulation of gene activity to support uncontrolled growth and survival. This regulation is controlled not only by genetic mutations, but also by epigenetic mechanisms, chemical and structural modifications to DNA and its associated proteins that determine which genes are turned on or off. Several therapies that target epigenetic regulators have shown promise, particularly in blood cancers. However, identifying which epigenetic mechanisms are most important in specific cancers remains challenging, especially because epigenetic regulators frequently work together as multi-protein complexes. In this study, we combine large-scale public datasets across many cancer lineages with computational modeling to systematically identify lineage-enriched epigenetic vulnerabilities. We found that certain epigenetic complexes are selectively important in specific cancer lineages. In melanoma, an aggressive skin cancer, we identified a previously unrecognized dependence on a protein complex that modifies chromatin at gene promoters. We show that disrupting this complex can impair gene programs that drive cell division and block cancer cell growth. Moreover, our study demonstrate how integrative computational approaches combined with experiments can uncover new targets for potential cancer therapy studies.

## Introduction

Epigenetic regulation plays a key role in controlling gene expression programs and maintaining stable cell identity during normal development and tissue homeostasis. This regulation is achieved through precisely coordinated mechanisms, including histone modifications, DNA methylation, chromatin remodeling, and higher-order genome organization. Together, these mechanisms regulate promoter accessibility, enhancer activity, and the engagement of transcriptional machinery with lineage-specific regulatory programs. Disruption of these epigenetic processes is a universal hallmark of cancer, leading to widespread transcriptional dysregulation, increased phenotypic heterogeneity, and enhanced cellular plasticity[1–3]. Importantly, such epigenetic dysregulation also creates selective dependencies on specific chromatin regulators and epigenetic mechanisms that become essential for tumor cell survival, proliferation, and adaptation[4].

Epigenetic vulnerabilities have been identified across multiple cancer types. In hematologic malignancies, recurrent alterations in chromatin regulators and epigenetic control mechanisms have informed the development of successful therapies, including FDA-approved DNA methyltransferase inhibitors, histone deacetylase inhibitors, as well as more recent agents targeting histone methyltransferases EZH2 and DOT1L, BET bromodomain proteins, and the histone demethylase LSD1[5–7]. Epigenetic vulnerabilities are also increasingly recognized in solid tumors, where cancer cells rely on chromatin-based mechanisms to maintain transcriptional programs associated with their oncogenic properties[4,8–10].

Even in melanoma, a cancer type associated with well-defined genetic driver alterations (including MAP kinase pathway-activating mutations in BRAF, NRAS, and NF1), these oncogenic drivers operate within a specific epigenetic and transcriptional landscape[11–17]. Melanoma cells exhibit remarkable phenotypic plasticity, transitioning between melanocytic, neural crest-like, and mesenchymal-like states, with these state transitions governed by epigenetic and transcriptional mechanisms[18–21]. Consistent with this, prior studies have identified multiple epigenetic vulnerabilities in melanoma through epigenomic profiling, functional perturbation, and epigenetic compound screening studies[11,22–25]. These efforts have implicated diverse chromatin regulators, including histone methyltransferases, demethylases, and chromatin remodelers, in controlling melanoma cell survival and transcriptional plasticity. However, these studies have largely focused on individual genes or regulators or specific functional classes of chromatin modifiers. Because chromatin regulators frequently assemble into multi-subunit complexes, epigenetic vulnerabilities ultimately depend on the complex activities rather than on single genes. Nevertheless, a systematic analysis that maps epigenetic complex-level dependencies across specific cancer lineages or defined differentiation states has not yet been performed. The availability of large-scale cancer dependency maps, integrated with transcriptomic and genomic datasets through multivariate computational modeling, provides a unique opportunity to address this gap.

In this paper, we present a systematic, multivariate analysis of epigenetic dependencies across >1,000 cancer cell lines and 42 major cancer types to identify epigenetic vulnerabilities that are selectively enriched in the melanoma lineage relative to other cancer lineages, as well as across distinct differentiation states within melanoma. Our analysis reveals multiple epigenetic complex dependencies, including complexes previously associated with recurrent genetic alterations or melanocyte lineage regulation, as well as a previously unrecognized vulnerability involving the H3K4 methyltransferase complex Set1C/COMPASS. This dependency is not restricted to a specific melanoma differentiation state. Instead, genetic perturbation of CXXC1, a core subunit of the complex, suggests that CXXC1-dependent melanoma cells rely on Set1C/COMPASS activity to maintain global H3K4 trimethylation (H3K4me3) and sustain cellular growth. Single-cell analysis reveals that loss of H3K4me3 following CXXC1 depletion is coupled to cell-cycle blockade in the top sensitive cell lines. Integrative analyses of cancer cell line transcriptomes and genome-wide gene dependencies link Set1C/COMPASS vulnerability to MYC- and E2F-driven transcriptional programs. Transcriptomic analysis following CXXC1 depletion confirms suppression of these programs in H3K4me3-responsive cell lines. Together, these findings identify Set1C/COMPASS as a melanoma-enriched epigenetic dependency that supports pro-growth transcriptional programs across diverse melanoma cell states, providing new insight into chromatin-mediated vulnerabilities in melanoma.

## Results

### Epigenetic dependency analysis of cancer cell lines reveals Set1C/COMPASS as a melanoma-enriched vulnerability

To systematically identify epigenetic dependencies that are differentially enriched in melanoma relative to other cancer lineages, we analyzed genome-wide DepMap gene-dependency scores from cell lines in the Cancer Cell Line Encyclopedia (CCLE)[26,27]. Our analysis included 42 cancer types, each represented by at least five CCLE cell lines. To evaluate dependencies at the level of chromatin-regulatory assemblies, we used a curated database of epigenetic regulators and their annotated protein complexes[28]. For each cell line, we calculated complex-level dependency scores by averaging gene-dependency values across the constituent subunits of each chromatin regulatory complex. Lineage-specific enrichment of complex dependencies was then assessed using a limma one-versus-rest model[29], comparing each cancer lineage against all remaining lineages. Given the exploratory nature of this analysis (see Methods for details), we applied a relatively permissive false discovery rate threshold (FDR ≤ 0.1) at this stage. Hierarchical clustering of these lineage-associated dependencies revealed groups of functionally related epigenetic complexes that were recurrently enriched across cancer lineages (Figure 1A). These included clusters of complexes involved in H3K4 methylation (Set1C/COMPASS, MLL1/MLL2, MLL3/MLL4, and Menin-MLL; highlighted in red), SWI/SNF-family ATP-dependent chromatin remodeling (PBAF, BAF, and ncBAF; highlighted in blue), ISWI-family chromatin remodelers (ACF, WICH/B-WICH, RSF, and NoRC; highlighted in purple), and SAGA-type histone acetyltransferases (STAGA/TFTC, PCAF, and SAGA; highlighted in orange). Cancer lineages were also segregated into clusters with distinct dependency profiles. Most prominently, hematopoietic malignancies (including mature B-cell neoplasms, acute myeloid leukemia, and B-cell acute lymphoblastic leukemia; highlighted in brown) clustered together due to elevated dependency on multiple chromatin regulatory complexes, consistent with previous reports[30–34]. Most solid tumors, however, showed selective dependencies on a limited number of complexes. Melanoma cell lines, specifically, exhibited enrichment for multiple epigenetic complex dependencies relative to other cancer types, including the H3K4 methyltransferase complex Set1C/COMPASS, the H3K9 methyltransferase complex G9a/GLP, the ATP-dependent chromatin remodeling complex NURF, the histone acetyltransferase complex HBO1, the ATRX-DAXX histone chaperone complex, the Elongator complex, and the BRCA1-A complex (Figure 1B). Several of these complexes have been functionally linked to melanoma biology; for example, inhibition of G9a/GLP reduces melanoma growth and induces apoptosis[35], and NURF regulates melanocyte lineage differentiation[36].

**Figure 1.**
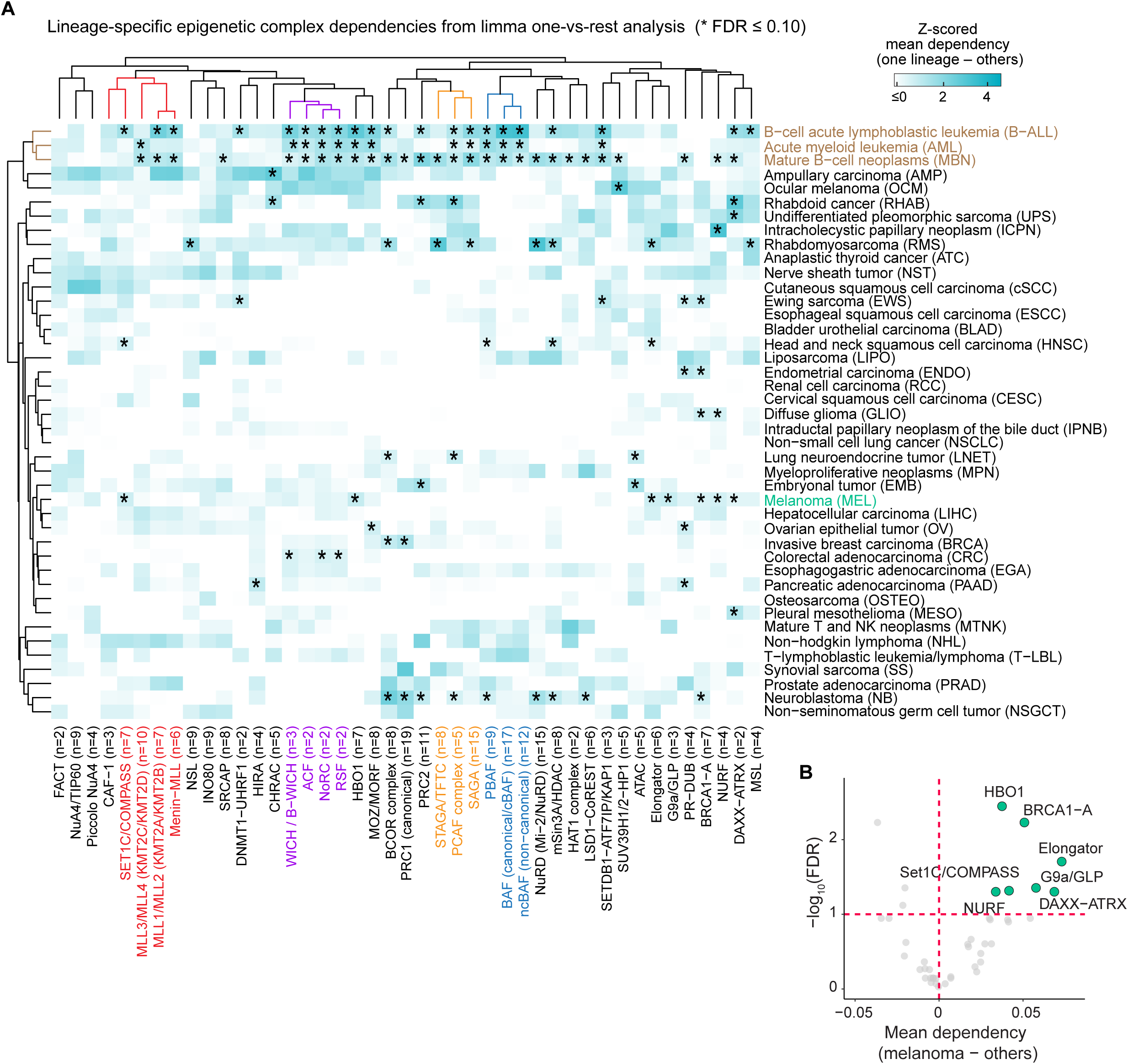
Analysis of epigenetic complex dependencies across cancer cell lines nominates Set1C/COMPASS dependency as an enriched vulnerability in melanoma. **(A)** Lineage-specific enrichment of epigenetic complex dependencies analyzed for CCLE cell lines across 42 cancer types using a limma one-versus-rest model. Data represent z-scored differential mean dependency (one lineage – all other lineages). Rows correspond to cancer types and columns to curated epigenetic complexes. Rows and columns were hierarchically clustered using Euclidean distance. * indicates candidate lineage-enriched complexes at an exploratory FDR ≤ 0.1 threshold. Column labels are colored to highlight functionally related clusters of epigenetic complexes: red, H3K4 methyltransferase complexes; blue, SWI/SNF-family chromatin remodelers; purple, ISWI-family chromatin remodelers; orange, SAGA-type histone acetyltransferase complexes. Row labels colored in brown represent several hematopoietic malignancies (acute leukemia/B-cell neoplasms) that were clustered together based on their shared epigenetic dependencies. The column label for the melanoma lineage is shown in green. **(B)** Enriched epigenetic complex dependencies observed for melanoma relative to all other lineages. Complexes with differential dependency > 0 and FDR ≤ 0.1 are highlighted.

In addition to the complex-level analysis, we took a complementary gene-level approach to identify statistically significant lineage-specific epigenetic dependencies by applying limma one-versus-rest models to 735 individual epigenetic regulator genes using a more stringent FDR ≤ 0.05. We identified 26 epigenetic regulator genes with significantly elevated dependency in melaonoma relative to all other lineages (mean difference in dependency > 0.1, FDR ≤ 0.05) (Figure 2A). We subjected this gene set to Enrichr over-representation analysis[37] (Figure 2B). In addition to two chromatin remodeling complexes (INO80-type and Swr1 complexes) and two histone acetyltransferase complexes (NuA4 and H4/H2A complexes), the H3K4 methyltransferase complex Set1C/COMPASS emerged again as a significantly enriched dependency, driven by the presence of its core subunits CXXC1 and SETD1B among the top 26 genes. Although only these two subunits met the significance threshold (FDR ≤ 0.05), two additional Set1C/COMPASS subunits (DPY30 and RBBP5) also showed higher mean dependency scores in melanoma cell lines compared to other lineages in the limma model (Figure 2C). Consistently, direct comparison of 67 melanoma cell lines versus 1,040 non-melanoma cell lines showed significantly higher dependency scores for both CXXC1 and SETD1B in melanoma (Figures 2D-E).

**Figure 2.**
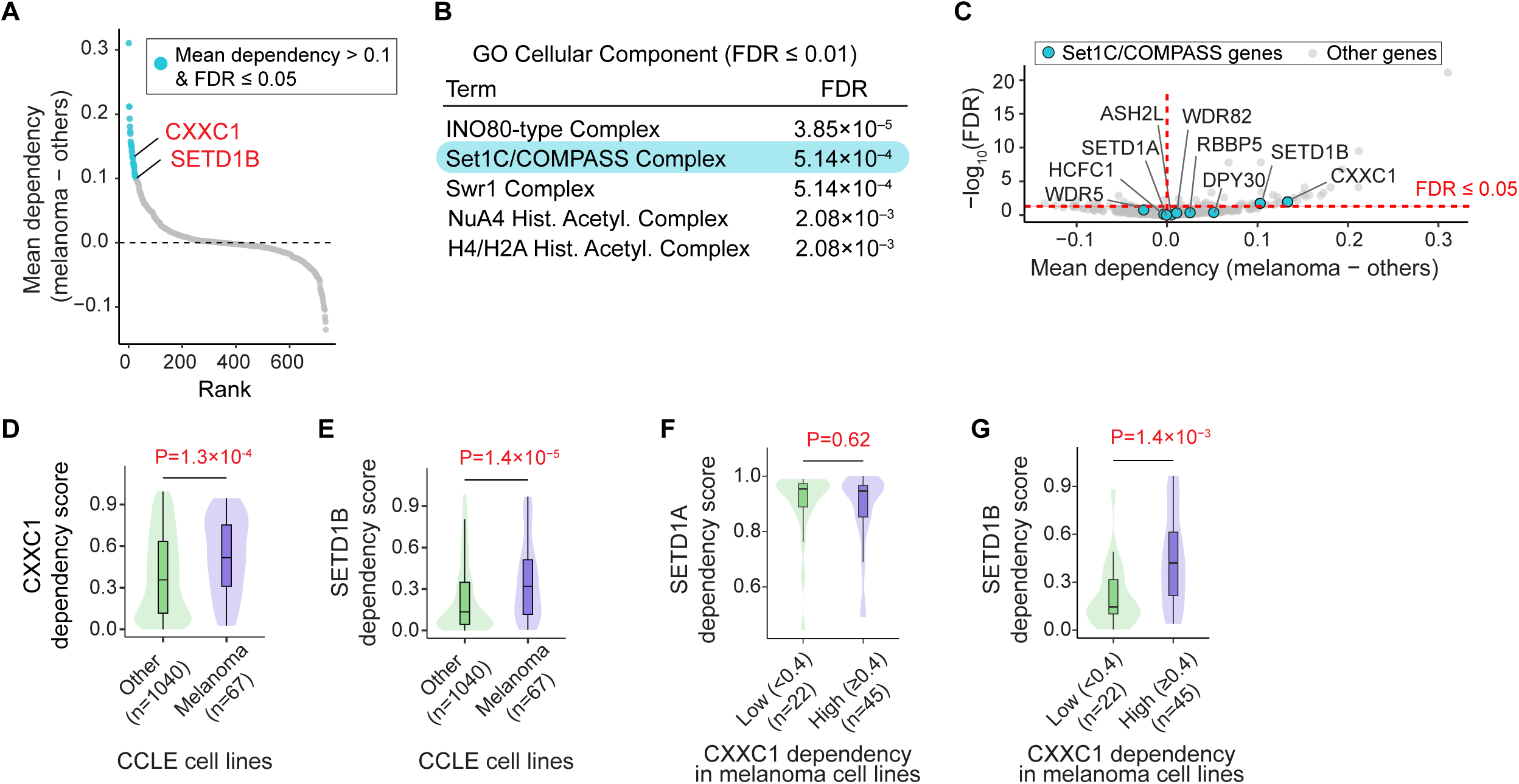
Analysis of epigenetic gene dependencies across cancer cell lines reveals significantly elevated dependency on the Set1C/COMPASS subunits CXXC1 and SETD1B in melanoma relative to other cancer lineages. **(A)** Epigenetic genes ranked based on their melanoma-enriched dependency. Top 26 genes (differential dependency > 0.1, FDR ≤ 0.05) are highlighted. CXXC1 and SETD1B (annotated in red) are among core subunits of the Set1C/COMPASS complex. **(B)** Over-representation analysis of melanoma-enriched epigenetic gene dependencies. The top 26 genes (differential dependency > 0.1, FDR ≤ 0.05) were submitted to Enrichr for over-representation analysis based on GO Cellular Component (C5: GO:CC). Shown are pathways corresponding to epigenetic complexes with FDR ≤ 0.01. Broad, non-specific compartments such as nucleus, intracellular membrane-bounded organelle, nuclear chromosome, nuclear lumen, chromosome, and nucleolus were excluded to focus on epigenetic complex terms. Entries are ranked by Benjamini-Hochberg adjusted P value (FDR). **(C)** Visualization of all 9 members of the Set1/COMPASS complex (ASH2L, CXXC1, DPY30, RBBP5, SETD1A, SETD1B, WDR5, WDR82, HCFC1) among melanoma-enriched epigenetic gene dependencies. **(D, E)** Comparison of CXXC1 and SETD1B dependency scores between melanoma cell lines and cell lines of all other lineages. **(F, G)** Comparison of SETD1A and SETD1B dependency scores between subgroups of melanoma cell lines stratified by CXXC1 dependency (CXXC1^Low^ vs CXXC1^High^; threshold = 0.4). Boxplots indicate the median and interquartile range, and the Wilcoxon rank-sum P values and group sample sizes (n) are shown for (D)-(G).

CXXC1 is a critical component of the Set1C/COMPASS complex that mediates recognition and binding to non-methylated CpG motifs via its CXXC domain, enabling establishment of H3K4 trimethylation (H3K4me3)[38]. H3K4 methylation within the Set1C/COMPASS complex can be catalyzed by either enzymes SETD1A or SETD1B. Despite their shared ability to methylate H3K4, SETD1A and SETD1B differ in both enzymatic properties and enzyme-independent functions[39–42]. Consistent with these differences, we observed uniformly high dependency on SETD1A across nearly all melanoma cell lines, independent of CXXC1 dependency (Figures 2F and S1A), whereas SETD1B dependency was heterogeneous and significantly correlated with CXXC1 dependency (Figures 2G and S1B). This correlated dependency on SETD1B and CXXC1 suggests that vulnerability in melanoma cell lines reflects their reliance on SETD1B-associated Set1C/COMPASS complex activity. Notably, Set1C/COMPASS has not previously been reported as a melanoma-associated epigenetic dependency. Furthermore, neither genetic mutations in Set1C/COMPASS subunits nor their expression levels correlated with CXXC1 dependency across melanoma cell lines (Figure S1C-E). We also examined whether global histone post-translational modification patterns correlated with CXXC1 dependency across CCLE melanoma cell lines. Among all modifications profiled, only H3K27ac1K36me0 showed a significant positive correlation with CXXC1 dependency (Figure S1F). However, this modification is not directly linked to the known function of the Set1C/COMPASS complex. We therefore next focused on exploring additional mechanisms that might be associated with Set1C/COMPASS dependency in heterogeneous melanoma populations.

### Melanoma-enriched Set1C/COMPASS dependency is not restricted to a specific melanoma differentiation state

The analysis above identified Set1C/COMPASS as a significant epigenetic vulnerability enriched in melanoma relative to other cancer lineages. However, melanoma tumors and cell lines exhibit substantial heterogeneity in differentiation state, which may itself be associated with distinct epigenetic vulnerabilities. We therefore sought to determine whether Set1C/COMPASS dependency in melanoma cell lines might be restricted to a specific differentiation state. Using RNA-sequencing (RNA-seq) data from 71 CCLE melanoma cell lines, we computed differentiation scores based on the z-scored expression of melanoma differentiation-state gene signatures (defined by Tsoi et al[19]), and applied a Gaussian mixture model to assign each cell line to either a melanocytic-like or an undifferentiated/neural crest-like class (Figures 3A and S2A). For the subset of 51 cell lines with available DepMap gene-dependency scores, we then applied multi-variate partial least-squares discriminant analysis (PLS-DA) to systematically identify epigenetic gene dependencies associated with each differentiation state.

**Figure 3.**
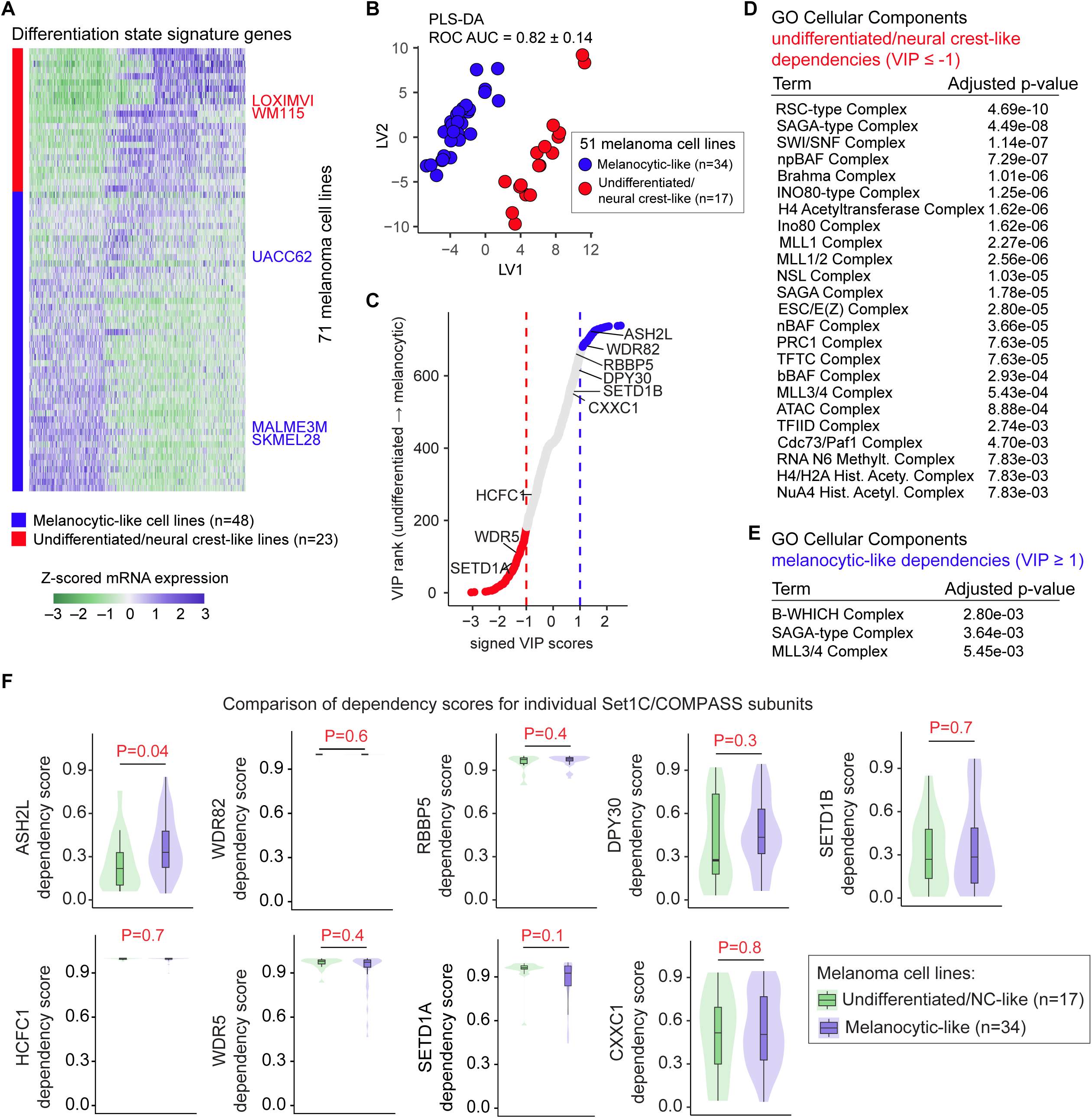
Melanoma-enriched Set1C/COMPASS dependency is not restricted to a specific melanoma differentiation state. **(A)** Classification of melanoma cell lines into melanocytic-like versus undifferentiated/neural crest-like groups based on the z-scored expression levels of Tsoi *et al* differentiation-state signature genes. Rows include 71 cell lines, and the position of representative cell lines from both differentiate state groups (used in follow-up studies) are highlighted in red and blue. **(B)** Partial least-squares discriminant analysis (PLS-DA) of epigenetic gene dependencies separates melanocytic-like cell lines from undifferentiated/neural crest-like cell lines. Data show PLS-DA latent variable 1 (LV1) versus LV2 scores. **(C)** Variable importance in projection (VIP) scores from the PLS-DA model ranked along the undifferentiated/neural crest-like → melanocytic axis. Each circle represents one epigenetic gene. Blue and red circles highlight genes with VIP ≥ 1 and VIP ≤ -1, representing significantly elevated dependencies in melanocytic-like and undifferentiated/neural crest-like lines, respectively. All nine Set1C/COMPASS subunits CXXC1, SETD1B, STED1A, DPY30, ASH2L, RBBP5, WDR82, WDR5 and HCFC1 are annotated. **(D, E)** EnrichR over-representation analysis by Gene Ontology (GO) Cellular Component showing enriched epigenetic genes with VIP ≤ -1 for undifferentiated/neural crest-like associated dependencies (D), and VIP ≥ 1 for melanocytic-associated dependencies (E). Broad, non-specific compartments such as nucleus, intracellular membrane-bounded organelle, nuclear lumen, nucleolus, and intracellular membrane-less organelle were excluded to focus on epigenetic complex terms. **(F)** Comparison of gene dependency scores for Set1C/COMPASS subunits between melanoma cell lines stratified by differentiation state class. P values indicate two-sided Wilcoxon rank-sum tests comparing dependency scores between classes, and n values indicate the number of cell lines per class (differentiation state).

PLS-DA separated the two differentiation states primarily along latent variable 1 (LV1), with a cross-validated ROC AUC of 0.82 ± 0.14 (Figures 3B and S2B), indicating statistically significant differences in epigenetic gene-dependency profiles between the two states. To identify genes driving this separation, we ranked epigenetic regulators using variable importance in projection (VIP) scores. Using a stringent |VIP| ≥ 1 threshold, we identified 78 genes as enriched vulnerabilities in melanocytic-like cell lines (VIP ≥ 1) and 178 genes as enriched vulnerabilities in undifferentiated/neural crest-like cell lines (VIP ≤ −1) (Figure 3C and Table S1). Enrichr over-representation analysis of these gene sets revealed several epigenetic complexes uniquely enriched (adjusted P value < 0.01) among undifferentiated/neural crest-like cell lines, including SWI/SNF family complexes, the INO80-type complex, and the NuA4 histone acetyltransferase complex (Figure 3D). Furthermore, components of the SAGA-type complex and the MLL3/4 histone methyltransferase complexes were associated with dependencies in both differentiation states (Figures 3D-E). However, Set1C/COMPASS dependency did not appear to be selectively enriched in either differentiation state. Consistent with this, comparison of dependency scores for individual Set1C/COMPASS subunits between melanoma cell lines from distinct differentiation states showed that only ASH2L dependency differed significantly between melanocytic-like and undifferentiated cell lines (Figure 3F). Thus, melanoma-enriched dependency on Set1C/COMPASS is not restricted to a specific melanoma differentiation state.

### CXXC1 depletion reduces H3K4me3 in CXXC1-dependent melanoma cell lines

To test whether dependency on CXXC1 in melanoma cells is associated with its role in supporting Set1C/COMPASS activity, we depleted CXXC1 using siRNA and assessed downstream effects on global H3K4me3, a readout of Set1C/COMPASS activity, as well as cellular growth and proliferation. To capture variability across CXXC1 dependency and melanoma differentiation states, we evaluated the effects of CXXC1 siRNA (si-CXXC1) relative to non-targeting control (si-NTC) across multiple melanoma cell lines spanning distinct differentiation states, including melanocytic lines MALME3M, UACC62, and SKMEL28 and undifferentiated lines LOXIMVI and WM115 (highlighted in Figure 3A). 96 h following siRNA treatment, we fixed the cells, counted them, and used iterative indirect immunofluorescence imaging (4i)[43] to quantify CXXC1 protein levels, H3K4me3, and the proliferation marker Ki-67 at the single-cell level (Figures 4A-B).

**Figure 4.**
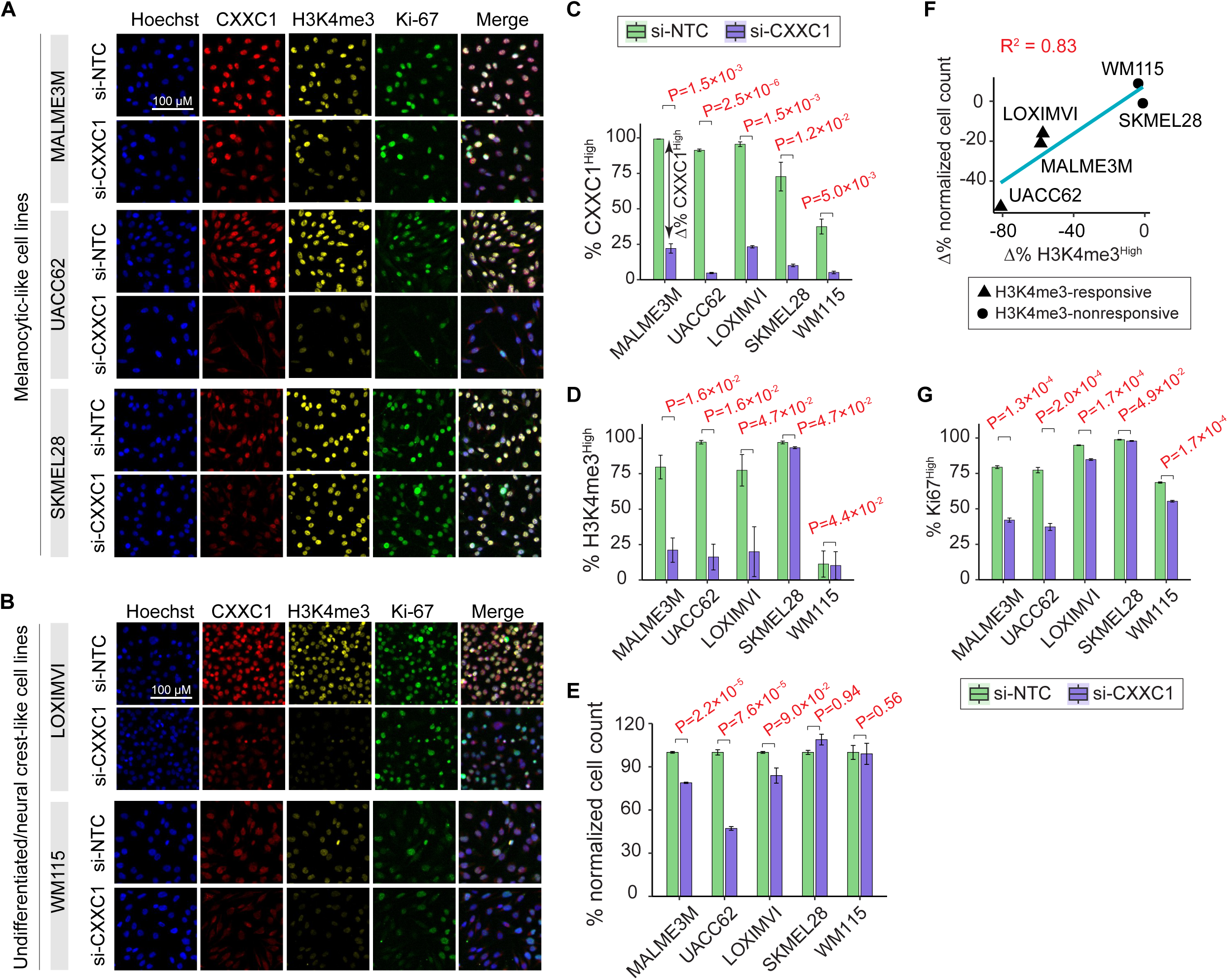
CXXC1 depletion reduces H3K4me3 in CXXC1-dependent melanoma cell lines. (A-B) Representative multiplexed immunofluorescence (4i) images of melanoma cell lines, treated with CXXC1 siRNA (si-CXXC1) and non-targeting control (si-NTC) for 96 h, followed by staining for CXXC1, H3K4me3, and Ki-67. Multiplexed images for three differentiated (melanocytic) cell lines (MALME3M, UACC62, SKMEL28) (A) and two undifferentiated/neural crest-like cell lines (LOXIMVI, WM115) (B) are shown. **(C-E)** Quantitative changes in the percentage of CXXC1^High^ cells (C), percentage of H3K4me3^High^ cells (D), and normalized cell count (E) following CXXC1 siRNA (si-CXXC1) treatment relative to non-targeting control (si-NTC) across five cell lines. Bars represent mean ± SEM from n = 3 biological replicates (wells). P values were derived from linear mixed-effects models with treatment as a fixed effect and well/replicate as a random effect, using one-sided tests adjusted for multiple comparisons across cell lines. **(F)** Relationship between percent change (Δ%) in H3K4me3^High^ cells and the corresponding percent change in normalized cell count following si-CXXC1 treatment across five melanoma cell lines. Percent changes were calculated relative to si-NTC. The solid line indicates the linear regression fit, and the coefficient of determination (R^2^) is shown in red. **(G)** Quantitative changes in the percentage of Ki-67^High^ cells following CXXC1 siRNA (si-CXXC1) treatment relative to non-targeting control (si-NTC) across five cell lines. Bars represent mean ± SEM from n = 3 biological replicates (wells). P values were derived as described for (C-E).

Across all cell lines, CXXC1 siRNA resulted in significant depletion of CXXC1 protein relative to non-targeting control (FDR < 0.05) (Figures 4C and S3A-B). Although the magnitude of knockdown varied across cell lines, every cell line exhibited more than a 30% reduction in the fraction of CXXC1^High^ cells following CXXC1 siRNA treatment relative to non-targeting control. Taking into account the differences in baseline CXXC1 expression, the estimated knockdown efficiency ranged from approximately 75% to 95%, indicating robust CXXC1 depletion in all five cell lines (Table S2A). However, only three cell lines, including MALME3M, UACC62, and LOXIMVI, exhibited a significant (FDR < 0.05) and greater than 30% reduction effect in the fraction of H3K4me3^High^ cells (Figures 4D and S3C-D and Table S2B). By comparison, SKMEL28 showed only a modest reduction (∼4%), whereas WM115 had a low baseline fraction of H3K4me3^High^ cells that did not change significantly (FDR = 0.4) following CXXC1 depletion. Based on these statistically different H3K4me3 responses following CXXC1 knockdown, we classified MALME3M, UACC62, and LOXIMVI as H3K4me3-responsive cell lines, and SKMEL28 and WM115 as H3K4me3-nonresponsive cell lines (Table S2B).

Next, using normalized cell count as a bulk readout of cellular growth 96 h following siRNA treatment, we found that two of the H3K4me3-responsive cell lines (MALME3M, UACC62) exhibited the strongest reductions in normalized cell count (∼21% and ∼53% reduction with effect size of ∼24 and ∼19, respectively) that were statistically significant (FDR < 0.05), whereas the H3K4me3-nonresponsive cell lines SKMEL28 and WM115 showed the weakest effect sizes (-1.8 and ∼0.1, respectively) that were not significant (FDR = ∼0.9 and ∼0.6, respectively; Figure 4E and Table S2C). For LOXIMVI (the third H3K4me3-responsive cell line), we observed only a modest reduction in normalized cell count (∼16%) with an effect size of 2.25 that did not meet the FDR cutoff for statistical significance (FDR = 0.09; Table S2C).

These results suggest a correspondence between the H3K4me3 response to CXXC1 depletion and CXXC1 dependency, as measured by the effect of CXXC1 knockdown on bulk cellular growth. Consistent with this, interaction tests confirmed that the effects of CXXC1 knockdown on both H3K4me3 and normalized cell count differed significantly (*P* ≈ 0.03 and *P* ≈ 0.002, respectively) between H3K4me3-responsive (MALME3M, UACC62, and LOXIMVI) and H3K4me3-nonresponsive (SKMEL28 and WM115) cell lines (Table S2D), supporting an association between H3K4me3 response and CXXC1 dependency. Accordingly, siRNA-induced changes in H3K4me3 were correlated with changes in normalized cell count across all five cell lines (Figure 4F).

We next asked whether the concurrent depletion of H3K4me3 and reduction in bulk cellular growth were accompanied by changes in the proliferative state of melanoma cells, as measured by Ki-67 positivity at the single-cell level. MALME3M and UACC62, the two cell lines exhibiting the greatest reductions in both H3K4me3 and normalized cell count following CXXC1 depletion, also showed the largest decreases (approximately 50%) in the fraction of Ki-67^High^ cells (Figures 4G and S3E-F). In contrast, LOXIMVI, as well as the two H3K4me3-nonresponsive cell lines SKMEL28 and WM115, exhibited only modest reductions (<20%) in the fraction of Ki-67^High^ cells (Figures 4G and S3E-F). Consistent with these data, MALME3M also showed a marked reduction in phosphorylation of Rb at Ser807/811 and Thr373, additional indicators of cell-cycle progression, whereas LOXIMVI exhibited a moderate effect and SKMEL28 showed no significant change in Rb phosphorylation following CXXC1 depletion (Figures S3G-H). Together, these results suggest that CXXC1 dependency in melanoma cells is associated with Set1C/COMPASS activity, as measured by global H3K4me3 levels. In H3K4me3-responsive cell lines, CXXC1 depletion results in loss of H3K4me3 and reduced cellular growth.

### Single-cell analysis confirms coupling of H3K4me3 loss to cell cycle blockade in the top sensitive cell lines

The population-level analyses above show that CXXC1 depletion reduces H3K4me3 globally and impairs proliferation in the most sensitive cell lines (MALME3M and UACC62), but they cannot determine whether these effects occur within the same cells or instead reflect independent effects in distinct subpopulations. To test whether loss of H3K4me3 and inhibition of proliferation occur within the same cells in these cell lines, we analyzed the covariance of these markers at the single-cell level using multiplexed imaging data.

In MALME3M and UACC62, most cells (∼80-90%) expressed high levels of both CXXC1 and H3K4me3 (CXXC1^High^/H3K4me3^High^) under baseline conditions (Figure 5A). Following CXXC1 depletion, the cell population underwent a redistribution toward a predominantly CXXC1^Low^/H3K4me3^Low^ state, comprising approximately 60-80% of cells (Figure 5A). These changes are consistent with CXXC1 depletion and loss of H3K4me3 being coupled at the single-cell level. Consistent with this coupling, siRNA-induced reduction in H3K4me3 was also associated with inhibition of proliferation marker Ki-67 at the single-cell level in both cell lines. In MALME3M and UACC62, cells shifted from a predominantly H3K4me3^High^/Ki-67^High^ state under baseline conditions to a largely H3K4me3^Low^/Ki-67^Low^ state following CXXC1 depletion (Figure 5B). Finally, single-cell covariance analysis of H3K4me3 with phosphorylated Rb and DNA content showed that loss of H3K4me3 following CXXC1 depletion was associated with a reduced fraction of cells in S and G2 phases and a corresponding increase in cells in G0/G1 (Figure 5C-D). Together, these data suggest that CXXC1 depletion in the most CXXC1-dependent melanoma cell lines leads to loss of H3K4me3, which is coupled to cell-cycle blockade in the G0/G1 phase.

**Figure 5.**
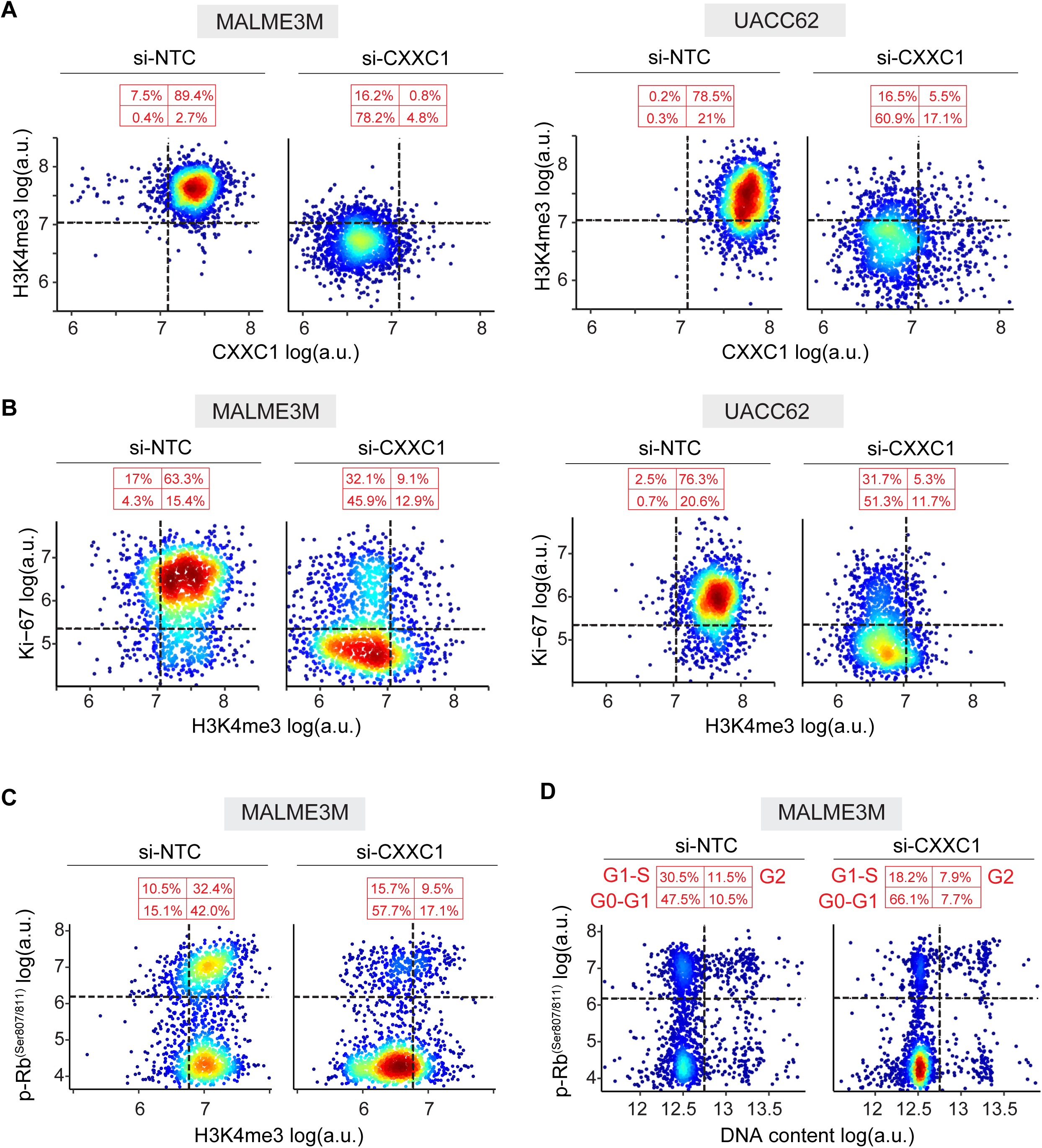
Single-cell analysis confirms coupling of H3K4me3 loss to cell cycle blockade in the top sensitive cell lines. **(A-B)** Single-cell covariate analysis of CXXC1 protein level versus H3K4me3 (A) and H3K4me3 versus Ki-67 (B) in MALME3M (left) and UACC62 (right) following 96 h treatment with CXXC1 siRNA (si-CXXC1) or non-targeting control (si-NTC). **(C, D)** Single-cell covariate analysis of H3K4me3 versus p-Rb^Ser807/811^ (C) and DNA content versus p-Rb^Ser807/811^ (D) in MALME3M cells following 96 h treatment with CXXC1 siRNA (si-CXXC1) or non-targeting control (si-NTC). 1,500 randomly sampled cells from each condition are visualized in each plot. Dotted horizontal and vertical lines denote Gaussian mixture model-derived thresholds used to gate low versus high fluorescence signal in cell populations. The average percentage of cells in each quadrant is indicated in red.

### Set1C/COMPASS dependency associates with MYC- and E2F-driven transcriptional programs in melanoma cell lines

As shown above, Set1C/COMPASS dependency in melanoma is not restricted to a specific differentiation state. We therefore asked whether it is associated with other transcriptional programs or biological pathways. To address this question, we analyzed 51 CCLE melanoma cell lines with matched RNA-seq data and genome-wide DepMap gene dependency scores. For each cell line, we calculated a Set1C/COMPASS dependency score by averaging the dependency scores for the core complex subunits CXXC1, SETD1B, ASH2L, DPY30, RBBP5, WDR5, and WDR82. We then performed pre-ranked gene set enrichment analysis (GSEA)[44] using MSigDB Hallmark gene sets, ranking genes according to the correlation between their expression and Set1C/COMPASS dependency across melanoma cell lines. To assess whether Set1C/COMPASS dependency was also associated with functional dependence on the same programs, we performed a parallel analysis using genome-wide gene-dependency data, ranking genes by the correlation between their dependency scores and the Set1C/COMPASS dependency score.

Several gene sets were negatively associated with Set1C/COMPASS dependency at both the expression and functional dependency levels, including epithelial-mesenchymal transition, KRAS signaling, and IL-2/STAT5 signaling (Figures 6A-B; green stars). In contrast, MYC targets, E2F targets, DNA repair, and oxidative phosphorylation were positively associated at both levels (Figures 6A-B; purple stars). Because Set1C/COMPASS maintains H3K4me3, a promoter-associated histone modification linked to open chromatin and transcriptional initiation, positively associated programs (such as MYC and E2F-linked transcriptional programs) are more likely to reflect direct effects of Set1C/COMPASS activity, whereas negatively associated programs may reflect more indirect relationships.

**Figure 6.**
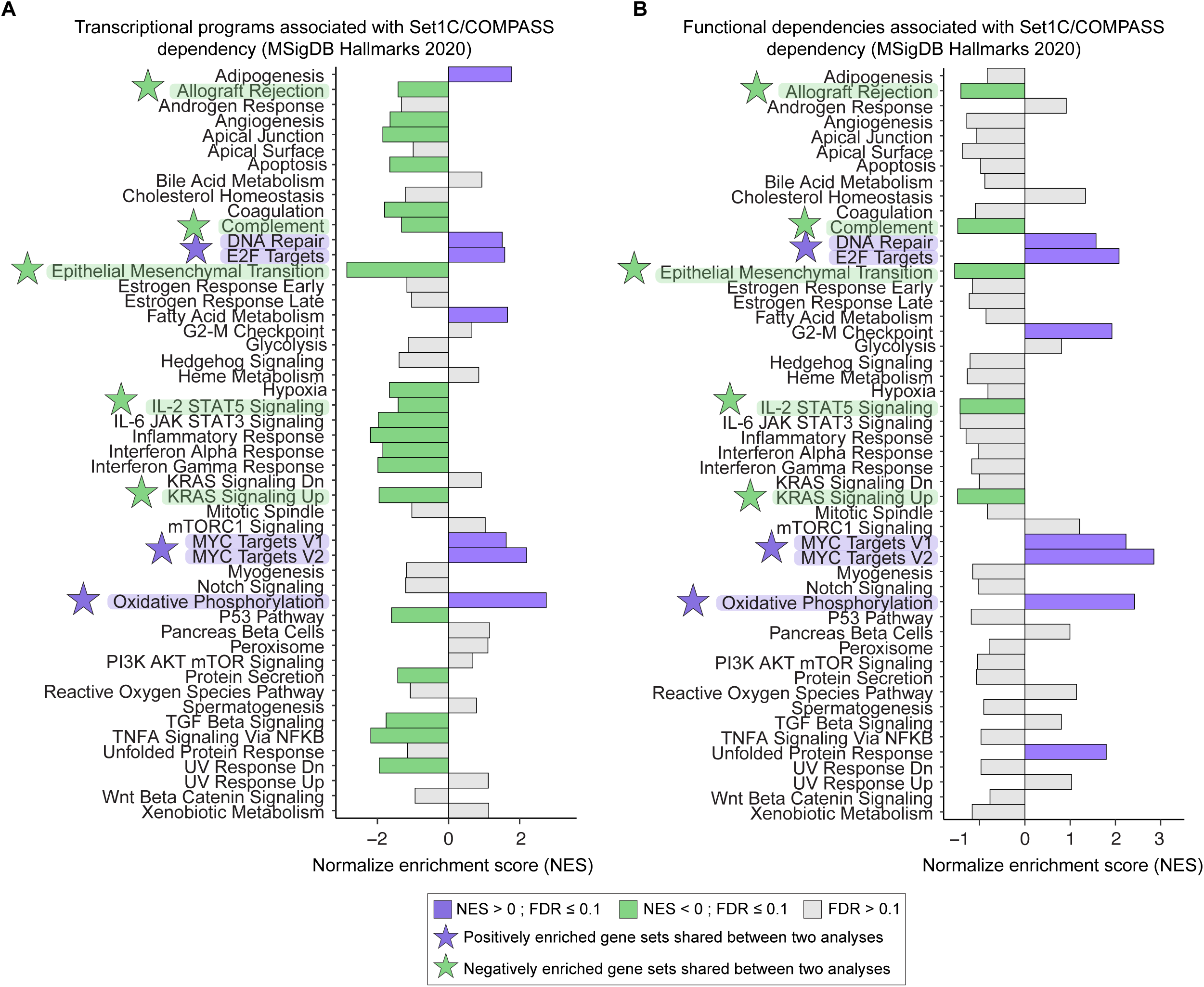
Set1C/COMPASS dependency associates with MYC- and E2F-driven transcriptional programs in melanoma cell lines. **(A, B)** Hallmark gene set enrichment analysis (fgsea; Spearman-ranked genes) across 51 melanoma cell lines with matched CCLE mRNA expression and DepMap gene dependency profiles. The phenotype was the mean Set1C/COMPASS dependency score (average of CXXC1, SETD1B, ASH2L, DPY30, RBBP5, WDR5, WDR82). Gene ranking was derived from mRNA expression (A) and from gene dependency scores (B). All Hallmark pathways are shown and annotated with FDR. Hallmark gene sets highlighted by purple and green stars denote positive and negative enrichment in both transcript-level (A) and gene dependency-level (B) analyses, respectively.

### CXXC1 depletion suppresses MYC- and E2F-linked transcription programs in H3K4me3-responsive melanoma cell lines

The analysis above shows that melanoma cell lines with greater Set1C/COMPASS dependency tend to exhibit both increased expression of and functional dependency on genes involved in MYC and E2F-linked transcriptional programs. This finding may suggest that inhibiting Set1C/COMPASS should attenuate MYC- and E2F-driven transcriptional programs in sensitive melanoma cells. We therefore next examined the transcriptional consequences of Set1C/COMPASS inhibition in our cell lines. To identify genes and pathways whose expression may be regulated by Set1C/COMPASS activity in melanoma cells, we performed RNA-seq analysis on four melanoma cell lines, including three H3K4me3-responsive lines (MALME3M, UACC62, and LOXIMVI) and one H3K4me3-nonresponsive line (SKMEL28) as a control. Cells were analyzed in biological triplicate 96 h following treatment with CXXC1 siRNA (si-CXXC1) or a non-targeting control (si-NTC). To systematically compare transcriptional responses across all samples, we first performed principal component analysis (PCA).

The first four principal components (PCs) together explained ∼84% of the total variance in the dataset (Figure 7A). Consistent with substantial genetic and transcriptional differences and differentiation-state heterogeneity among four cell lines, the first three PCs primarily captured cell line-specific variation independent of treatment (Figure S4A). In contrast, PC4 significantly separated si-NTC and si-CXXC1 samples in MALME3M, UACC62, and LOXIMVI, whereas no significant separation was observed in SKMEL28 (Figure 7B). Notably, all four cell lines exhibited significant CXXC1 transcript depletion following siRNA treatment (Figure S4B); PC4 therefore appeared to capture the transcriptional response to CXXC1 depletion specifically in H3K4me3-responsive cell lines.

**Figure 7.**
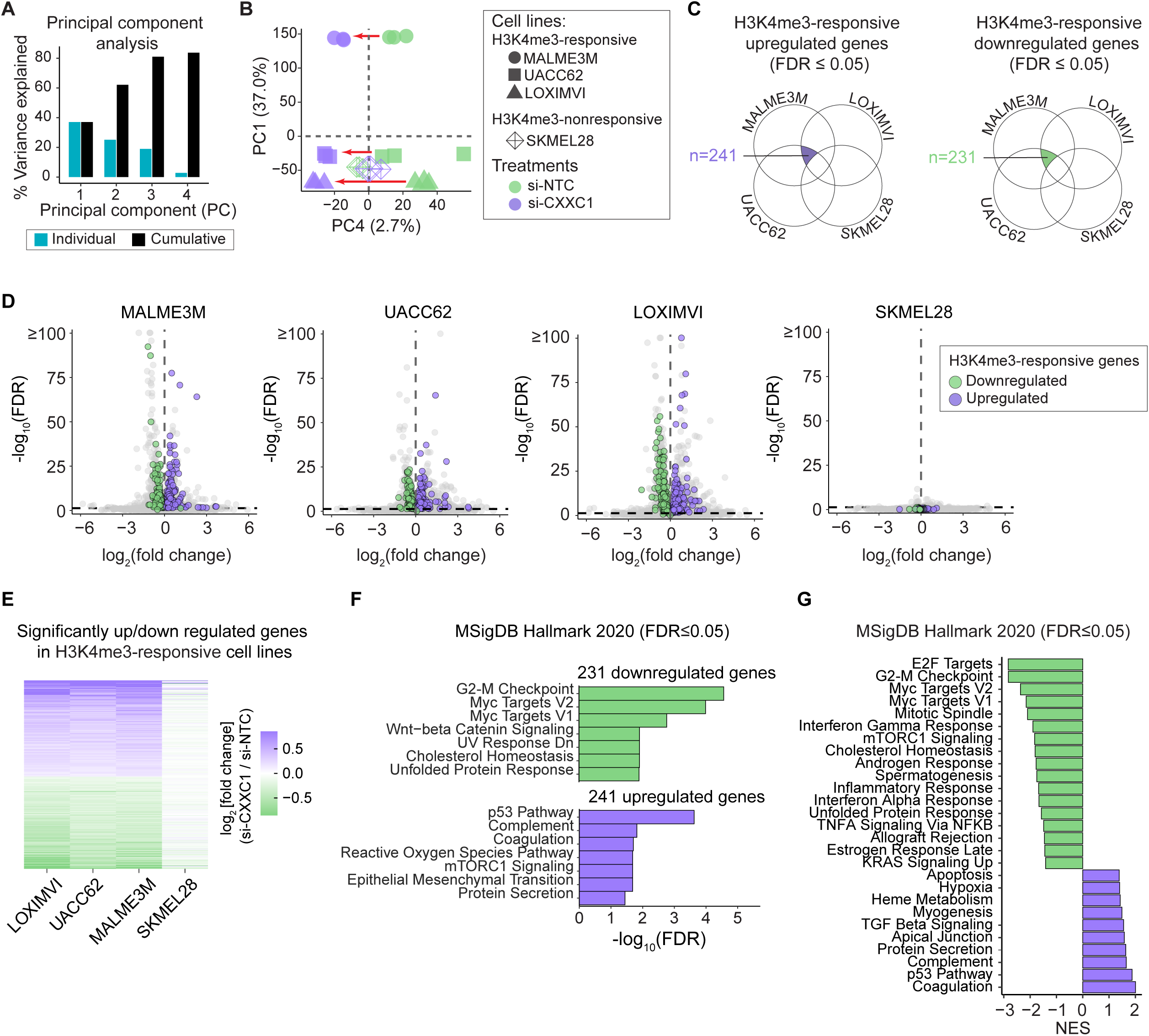
CXXC1 depletion suppresses MYC- and E2F-linked transcription programs in H3K4me3-responsive melanoma cell lines. **(A)** PCA of gene expression variations across 12 samples, representing three H3K4me3-responsive cell lines (MALME3M, UACC62, LOXIMVI) and one H3K4me3-nonresponsive cell line (SKMEL28) following 96 h treatment with CXXC1 siRNA (si-CXXC1) or the non-targeting control (si-NTC), each tested in three replicates. Percent variance captured by the first four PCs are shown. **(B)** PC1 versus PC4 scores capturing treatment-specific variations in gene expression in H3K4me3-responsive cell lines. Arrows show significant changes of scores by treatment in H3K4me3-responsive cell lines. **(C)** Schematic of the filtering strategy used to define consensus gene sets across H3K4me3-responsive melanoma cell lines. Differentially expressed genes (si-CXXC1 vs si-NTC; DESeq2 Wald test) were first intersected across LOXIMVI, MALME3M, and UACC62 within each direction (upregulated or downregulated). Genes that were also significantly altered in SKMEL28 in the same direction were excluded, yielding a final set of genes consistently changed in H3K4me3-responsive lines but not shared with the H3K4me3-nonresponsive reference. **(D)** Volcano plots for each melanoma cell line showing log_2_FC (si-CXXC1 vs si-NTC) versus -log_10_(FDR). The dashed horizontal line marks the FDR threshold (FDR = 0.05). Genes meeting the consensus criteria in H3K4me3-responsive lines are highlighted (purple, consensus up; green, consensus down). **(E)** Aggregated list of H3K4me3-responsive genes and their expression changes across cell lines. **(F)** Enrichr over-representation analysis of the consensus (H3K4me3-responsive) upregulated and downregulated gene sets using MSigDB Hallmark gene sets. Bars show the top enriched pathways passing FDR ≤ 0.05, ranked by adjusted P value. **(G)** Gene set enrichment analysis (GSEA) using a consensus ranked list derived from DESeq2 Wald statistics for the si-CXXC1 versus si-NTC contrast. Bars show normalized enrichment scores (NES), with positive NES indicating gene sets enriched toward genes upregulated in si-CXXC1 and negative NES indicating enrichment toward genes upregulated in si-NTC; only gene sets with FDR ≤ 0.05 are shown.

To identify genes and pathways underlying this selective transcriptional response, we applied two complementary approaches. First, we performed differential gene expression analysis for each cell line comparing si-CXXC1 versus si-NTC conditions and identified genes that changed in the same direction (upregulated or downregulated) across all three H3K4me3-responsive cell lines, but not in SKMEL28 (Figure 7C). This analysis yielded 231 genes that were consistently downregulated and 241 genes that were consistently upregulated in MALME3M, UACC62, and LOXIMVI following CXXC1 depletion, but did not show consistent changes in SKMEL28 (Figures 7D-E). Over-representation analysis of the downregulated genes revealed significant enrichment of cell-cycle and proliferation-associated Hallmark gene sets, with G2-M checkpoint and MYC targets emerging as the top categories (Figure 7F, top panel). In contrast, the p53 pathway was the most significantly enriched among upregulated gene sets (Figure 7F, bottom panel).

As a complementary approach that does not rely on hard thresholds for differential expression, we next assessed coordinated pathway-level changes using gene set enrichment analysis (GSEA). To do so, we constructed a merged gene ranking based on the mean DESeq2[45] Wald statistic across the three H3K4me3-responsive cell lines and performed pre-ranked GSEA using Hallmark gene sets. This analysis again identified E2F targets, G2-M checkpoint, and MYC targets among the most strongly enriched gene sets following CXXC1 depletion, whereas the p53 pathway was among the most negatively enriched (Figure 7G). Using the same merged ranking, GSEA with KEGG pathways or Gene Ontology Biological Process terms further confirmed broad downregulation of pathways and processes related to cell cycle progression, DNA replication and cell division (Figures S4C-D).

These data suggest that CXXC1 depletion induces a transcriptional response specifically in H3K4me3-responsive melanoma cell lines, characterized by suppression of MYC- and E2F-driven cell-cycle programs. This is consistent with the CCLE-based analyses described above (Figure 6), which linked Set1C/COMPASS dependency to MYC- and E2F-driven transcriptional programs in melanoma. We therefore asked whether this association is melanoma-specific or represents a more general feature of Set1C/COMPASS dependency across cancer lineages. Although our large-scale pan-cancer analysis (Figure 1A) identified significant enrichment of Set1C/COMPASS dependency in only a few cancer types, including melanoma (MEL), B-cell acute lymphoblastic leukemia (B-ALL), and head and neck squamous cell carcinoma (HNSC), many other lineages contained subsets of cell lines with elevated Set1C/COMPASS dependency. Moreover, a recent study showed that SETD1B-dependent H3K4 trimethylation is required to maintain MYC expression and cellular proliferation in acute myeloid leukemia (AML) [46]. We therefore extended the analysis presented in Figure 6 to all cancer lineages represented by at least five cell lines. For each lineage, we performed lineage-specific pre-ranked GSEA, ranking genes according to the correlation between their dependency scores and the Set1C/COMPASS dependency score across cell lines within that lineage. Remarkably, this association extended beyond melanoma, with most cancer lineages, including B-ALL, HNSC, and AML, exhibiting significant enrichment of MYC and E2F target gene dependencies among genes positively correlated with Set1C/COMPASS dependency (Figure S5). This association was particularly strong in HNSC, another cancer lineage showing significant enrichment of Set1C/COMPASS dependency in our pan-cancer analysis. Together, these data suggest that although the prevalence of Set1C/COMPASS dependency varies across cancer lineages and is enriched in only a subset of cancers, its functional association with MYC- and E2F-driven transcriptional programs may be broadly conserved. However, the molecular determinants of Set1C/COMPASS dependency, and the basis for its differential enrichment across cancer lineages, remain to be elucidated.

## Discussion

Epigenetic dysregulation is a feature of many cancers and creates vulnerabilities that arise due to an increased reliance on chromatin-based mechanisms that sustain malignant transcriptional states. However, many chromatin regulators are broadly required for cellular viability, while others exert context-dependent functions that vary across tissue lineages, differentiation states, and transcriptional programs. Furthermore, epigenetic vulnerabilities are often difficult to discern using single-gene perturbations, particularly when dependencies arise from cooperative activity within multi-subunit chromatin complexes[47]. In this study, we integrated large-scale gene dependency maps for human cancer cell lines with curated epigenetic complex annotations to perform a systematic, multivariate analysis of epigenetic dependencies enriched across cancer lineages, thereby providing a way to relate specific gene vulnerabilities to underlying lineage-specific chromatin mechanisms. Focusing on melanoma, our analysis identified multiple epigenetic complex dependencies, including complexes previously linked to recurrent genetic alterations in melanoma[48] or to melanocyte lineage differentiation[36], as well as a previously unrecognized vulnerability involving the H3K4 methyltransferase complex Set1C/COMPASS.

The Set1C/COMPASS complexes containing either the SETD1A or SETD1B catalytic subunits represent two of the six members of the larger COMPASS family of H3K4 methyltransferases, which also includes the MLL1, MLL2, MLL3, and MLL4 complexes[38]. Although all COMPASS family members catalyze H3K4 methylation and share a common core of subunits (WDR5, RBBP5, ASH2L, and DPY30), they assemble into distinct complexes with different genomic targeting and functional roles[49]. MLL1 and MLL2 primarily mediate locus-specific H3K4me3 at select promoters[50–52], MLL3 and MLL4 function mainly at enhancers as H3K4 monomethyltransferases[53,54], whereas SETD1A and SETD1B complexes are considered the principal contributors to bulk promoter-associated H3K4 trimethylation[55–57]. A distinguishing feature of SETD1A/SETD1B complexes is the CXXC1 subunit, which recruits the complex to non-methylated CpG islands to facilitate H3K4me3 deposition[38]. Therefore, the selective enrichment of dependency on CXXC1 and SETD1B in melanoma cell lines relative to other cancer lineages suggests a vulnerability linked either to a specific function of the SETD1B-containing COMPASS complex or to a specific biological context in which this complex becomes important in melanoma. Notably, this dependency was not explained by genetic alterations in, or expression of, Set1C/COMPASS subunits or other COMPASS family members and was not restricted to a specific melanoma differentiation state. Defining the molecular basis of this epigenetic vulnerability, and the sources of heterogeneity in Set1C/COMPASS dependency across melanoma populations, remain important directions for future investigation.

Our CXXC1 perturbation experiments suggest that Set1C/COMPASS dependency in melanoma cell lines is associated with a requirement to maintain H3K4 trimethylation to sustain MYC- and E2F-driven transcriptional programs and cell-cycle progression. This model is consistent with a recent study in leukemia showing that SETD1B-dependent H3K4 methylation supports MYC expression and cytokine-independent cellular proliferation[46]. However, additional mechanistic studies will be needed to further establish the causal relationship between CXXC1 expression, Set1C/COMPASS activity, and regulation of MYC- and E2F-driven transcriptional programs. For example, rescue experiments using wild-type CXXC1 or complementary Set1C/COMPASS subunits (e.g., SETD1B) could further validate the specificity of the observed growth defect and exclude potential off-target effects of CXXC1 siRNA. In addition, the mechanism linking Set1C/COMPASS activity to MYC- and E2F-driven transcription remains to be defined. Our current data cannot distinguish whether Set1C/COMPASS directly regulates MYC and E2F target genes through H3K4me3 deposition at their promoters or acts through indirect mechanisms. Future chromatin occupancy studies, such as ChIP-seq or CUT&RUN for CXXC1 and H3K4me3, will therefore be important to determine whether these transcriptional programs are direct targets of Set1C/COMPASS activity in melanoma cells.

Although the prevalence of Set1C/COMPASS dependency varies substantially across cancer lineages, our pan-cancer analysis of CCLE cell lines suggests that its functional association with MYC-and E2F-driven transcriptional programs may not be specific to melanoma, but instead broadly conserved across cancer lineages. This observation may point to a potential mechanism linking Set1C/COMPASS-mediated chromatin regulation to MYC- and E2F-dependent transcriptional programs across diverse cancer types. However, the molecular determinants of Set1C/COMPASS dependency remain unclear. In particular, future studies are needed to determine what predicts this dependency in tumor cells and why its prevalence differs across cancer lineages.

In melanoma, MYC activity is frequently deregulated through chromosome 8q24 amplification or activating mutations in the RAS/RAF/MAPK pathway, contributing to aggressive disease behavior and resistance to targeted therapies[58,59]. Thus, Set1C/COMPASS vulnerability in melanoma may arise as a mechanism that supports and stabilizes MYC-driven proliferative programs initiated by these genetic alterations. Notably, a clinically advanced MYC inhibitor has recently shown efficacy in melanoma cell lines and mouse models, suppressing tumor growth and reducing metastatic capacity by attenuating MYC-dependent transcriptional programs[60,61]. Our findings suggest that targeting Set1C/COMPASS may represent an additional pathway to disrupt these oncogenic transcriptional programs. However, further studies will be needed to evaluate the *in vivo* relevance of this dependency and to explore the therapeutic potential of pharmacologic inhibition of Set1C/COMPASS.

## Materials and Methods

### Cell culture

BRAF-mutant melanoma cell lines used in this study include: LOXIMVI, MALME3M, SKMEL28, UACC62 and WM115. All cell lines have been subjected to reconfirmation by short tandem repeat (STR) profiling by ATCC and mycoplasma testing by MycoAlert PLUS Mycoplasma Detection Kit. SKMEL28, and WM115 cells were grown in DMEM/F12 supplemented with 1% sodium pyruvate and 5% FBS. MALME3M, LOXIMVI, and UACC62 cells were grown in RPMI 1640 supplemented with 1% sodium pyruvate and 5% FBS. 100 U/mL Penicillin-Streptomycin (10,000 U/mL) and 0.5 mg/mL Plasmocin Prophylactic were present in all cell cultures. Cells were grown at 37 °C with 5% CO2 in a humidified incubator.

### Reagents

The following primary monoclonal antibodies (mAbs) with specified animal sources, catalog numbers and dilution ratios, were used in immunofluorescence (IF) staining: CXXC1 (CGBP) (rabbit mAb, Abcam, catalog no. ab198977, IF, 1:800), Ki-67 (AB_2797703; mouse mAb, Cell Signaling Technology, catalog no. 9449, IF, 1:800), H3K4me3 (C42D8) (AB_2616028; rabbit mAb, Cell Signaling Technology, catalog no. 9751, IF, 1:1600), p-Rb^S807/811^ (D20B12) (AB_11178658; rabbit mAb, Cell Signaling Technology, catalog no. 8516, IF, 1:1000), p-Rb^T373^ (rabbit mAb, Abcam, catalog no. ab52975, IF, 1:1000). The following secondary antibodies with specified sources and catalog numbers were used at 1:2,000 dilution for IF assays: anti-mouse Alexa Fluor 568 (AB_11180865; Thermo Fisher Scientific, catalog no. A10037) and anti-rabbit Alexa Fluor 647 (AB_2536183; Thermo Fisher Scientific, catalog no. A31573).

### Iterative indirect immunofluorescence imaging (4i)

4i images were acquired using a cell culture-based 4i protocol previously described in detail[43]. For experiments assessing the effects of CXXC1 knockdown, cells were seeded in 100 µl per well in Corning 96-well plates. LOXIMVI, MALME3M, SKMEL28, UACC62, and WM115 cells were plated at 1,600, 2,000, 1,300, 2,000 and 3,500 cells per well, respectively, with three biological replicates per condition. Cells were imaged using a 10x air objective on the Operetta CLS High-Content Imaging System (Revvity).

### 4i image analysis

Quantification of 4i immunofluorescence data was performed using an established image-processing workflow[21,43]. Raw images were first background-subtracted in ImageJ (v2.17.0) using the rolling-ball subtraction algorithm (radius = 50 pixels). Background-subtracted Hoechst images were subsequently used to align imaging rounds in CellProfiler (v4.2.8) with the Align module based on normalized cross-correlation. Nuclear segmentation was performed on the Hoechst channel using parameters optimized to best capture cell line-specific nuclear morphology. For knockdown experiments in LOXIMVI, MALME3M, UACC62, SKMEL28, and WM115, nuclei were identified using the IdentifyPrimaryObjects module with Minimum Cross-Entropy thresholding (smoothing scale = 2; correction factor = 1.7). Nuclei were tracked across imaging rounds using the TrackObjects module with the Follow Neighbors method. Mean nuclear intensity and/or integrated intensity values were extracted for all markers and used for downstream quantitative analyses. Data organization, summary calculations, and statistical analyses were performed in R.

### CXXC1 knockdown by siRNA (SMARTpool)

CXXC1 was depleted using an ON-TARGETplus siRNA SMARTpool targeting human CXXC1 (Horizon Discovery Biosciences; catalog no. L-014923-01-0005) and compared to an ON-TARGETplus Non-targeting siRNA #1 control (catalog no. D-001810-01-05). siRNA transfections were performed using DharmaFECT transfection reagents (Horizon Discovery), including DharmaFECT 1 (catalog no. T-2001-02), DharmaFECT 2 (catalog no. T-2002-01), and DharmaFECT 3 (catalog no. T-2003-01), according to the manufacturer’s instructions.

For immunofluorescence (IF) analysis of samples following knockdown, cells were seeded in 96-well plates at the densities specified below and allowed to attach for 24 hours before siRNA-DharmaFECT complexes were added. After 96 hours of knockdown, plates were fixed in 4% paraformaldehyde (PFA) before downstream 4i assays. Optimized conditions per cell line for the 96-well IF experiments were as follows (media; DharmaFECT reagent; DharmaFECT volume per well; cells seeded per well in 100 µL): LOXIMVI (RPMI-1640; DharmaFECT 3; 0.0500 µL; 1,600 cells); MALME3M (RPMI-1640; DharmaFECT 2; 0.0625 µL; 2,000 cells); SKMEL28 (DMEM/F-12; DharmaFECT 1; 0.1250 µL; 1,300 cells); UACC62 (RPMI-1640; DharmaFECT 1; 0.0625 µL; 2,000 cells); WM115 (DMEM/F-12; DharmaFECT 1; 0.0625 µL; 3,500 cells).

For bulk RNA-seq experiments, the same transfection protocol was applied in 6-well plates, with cell line-specific seeding densities and DharmaFECT volumes adjusted accordingly. After 96 hours, cells were lysed using the RNeasy kit (Qiagen) for RNA extraction.

### RNA extraction and library preparation

LOXIMVI, MALME3M, UACC62, and SKMEL28 cells were seeded in 6-well plates in biological triplicate and allowed to attach for 24 h prior to siRNA transfection. Cells were then treated with ON-TARGETplus siRNA SMARTpool targeting human CXXC1 or an ON-TARGETplus non-targeting control and cultured for 96 h post-transfection (transfection conditions and reagents are described above). At harvest, cells were washed once with PBS and lysed directly in the well using Buffer RLT (Qiagen). Lysates were homogenized using QIAshredder columns (Qiagen, catalog no. 79654), and total RNA was extracted using the RNeasy Mini Kit (Qiagen, catalog no. 74104) following the manufacturer’s protocol, including on-column DNase digestion (Qiagen RNase-Free DNase Set, catalog no. 79254). RNA integrity was assessed using the Agilent TapeStation (RNA ScreenTape), and all samples used for sequencing had RINe values > 9. Bulk RNA sequencing libraries were prepared with assistance from the University of Virginia Genome Analysis and Technology Core (RRID: SCR_018883). Poly(A)+ RNA was isolated using the NEB mRNA Isolation Module, followed by library preparation using the NEBNext UltraExpress mRNA Library Prep Kit, according to the manufacturers’ instructions. Libraries were sequenced on an Illumina NextSeq 2000 platform using a P3-100 sequencing kit. Raw FASTQ files were subjected to initial quality control using FastQC. Adapter sequences and low-quality bases were removed from paired-end reads using Trim Galore, with post-trimming quality assessment performed using FastQC. Trimmed reads were aligned to the human reference genome (GRCh38) using STAR[62], with genome indices generated from the GENCODE v41 primary assembly FASTA and annotation files. STAR was run in paired-end mode to generate coordinate-sorted BAM files as well as transcriptome-aligned BAMs for downstream quantification. Alignment quality and read distribution metrics were assessed using Qualimap RNA-seq. Transcript-and gene-level expression quantification was performed using RSEM[63] with a reference prepared from the same GENCODE v41 annotation, yielding expected counts and TPM values. Gene-level expected counts were compiled across all samples into a single expression matrix for downstream analysis.

### Analysis of gene expression data following CXXC1 knockdown

Principal component analysis (PCA) was performed on a gene expression matrix in which rows corresponded to genes, and columns corresponded to samples, using log2-transformed TPM+1 values. Prior to PCA, expression values were standardized on a per-gene basis by calculating row-wise z-scores across all samples (each gene was mean-centered across samples and divided by its standard deviation). The standardized matrix was then transposed to a samples × genes format for PCA input. PCA was computed in R using the prcomp function without additional scaling. The percent variance explained by each principal component was calculated from the component eigenvalues, and cumulative variance explained was computed as the running sum across components.

Differential expression analysis was then performed using DESeq2 on raw gene-level counts[45]. Gene identifiers were parsed from the count matrix row names to extract Ensembl gene IDs and HGNC symbols and counts mapping to the same Ensembl ID were collapsed by summation. For each cell line, differential expression between si-CXXC1 and si-NTC was tested using DESeq2 with Wald tests. Low-abundance genes were filtered prior to model fitting (row sum of counts across all samples ≥ 10). Adjusted P values were computed using the Benjamini-Hochberg method. For each H3K4me3-responsive melanoma cell line (MALME3M, UACC62, LOXIMVI), genes were classified as significantly upregulated or downregulated if FDR ≤ 0.05 and the estimated log2 fold change was > 0 or < 0, respectively.

To interpret transcriptional changes at the pathway level, two complementary enrichment strategies were applied: a cutoff-based consensus gene set approach followed by over-representation analysis, and a rank-based gene set enrichment approach using genome-wide statistics. For the cutoff-based approach, a consensus H3K4me3-responsive line response signature was defined by intersecting genes that were significantly upregulated or downregulated in all three H3K4me3-responsive lines with a consistent direction of change. To compare against an H3K4me3-nonresponsive reference line (SKMEL28), genes that were also significantly altered in SKMEL28 in the same direction were excluded, while genes that were either non-significant in SKMEL28 or changed in the opposite direction were retained. Functional enrichment of consensus upregulated and downregulated gene sets was evaluated using Enrichr by submitting HGNC gene symbols as input[37,64]. Enrichment was run separately for upregulated and downregulated consensus gene lists using curated pathway and gene set libraries. Enriched terms were interpreted using Enrichr’s multiple-testing corrected adjusted P values (FDR), and significance was defined as FDR ≤ 0.05.

In parallel, gene set enrichment analysis (GSEA) was performed using fgsea to identify coordinated pathway-level shifts without applying a hard differential expression cutoff[44]. For each H3K4me3-responsive cell line, genes were ranked by the DESeq2 Wald statistic from the si-CXXC1 versus si-NTC comparison, which captures both the direction and magnitude of differential expression. A merged H3K4me3-responsive line ranking was constructed by intersecting genes present across all three H3K4me3-responsive lines and computing the mean Wald statistic for each gene across lines, generating a single consensus ranked list (positive values indicate consistent upregulation following si-CXXC1; negative values indicate consistent downregulation). Pathway gene sets were obtained from MSigDB (Hallmark), GO:BP, and KEGG, and enrichment analysis was performed using fgsea. Gene set size constraints were enforced (minimum size 15 and maximum size 500 for all collections). Multiple-testing correction was performed using the Benjamini-Hochberg method. Enrichment results were summarized using normalized enrichment scores (NES), and pathways were considered enriched at FDR ≤ 0.05.

### Classification of single cells based on high or low expression of protein markers

To classify single cells into low versus high states for each immunofluorescence (4i) protein marker, per-cell signal intensities were log-transformed and pooled across all cell lines and treatment conditions. A two-component Gaussian mixture model was then fit to the resulting one-dimensional log-intensity distribution, and posterior component membership was used to assign each cell to either the low or high component. The classification threshold was defined as the analytic decision boundary at the intersection of the two fitted component density curves, accounting for mixture proportions. This single threshold was then applied uniformly across all cell lines and treatment conditions, with classification performed independently of cell line identity or treatment condition. Notably, this global pooling approach uses the full range of variation observed for each protein marker across the experiment to establish a single, stable classification threshold.

### Differentiation state classification of melanoma cell lines

Differentiation-state classes were assigned using a melanoma differentiation gene-signature table curated by Tsoi et al., in which each signature gene has been annotated to one of seven ordered transcriptional states (melanocytic, transitory-melanocytic, transitory, neural crest-like-transitory, neural crest-like, undifferentiated-neural crest-like, and undifferentiated)[19]. Gene expression was quantified as log-transformed TPM values (log2(TPM+1)) across a panel of 71 melanoma cell lines. To enable comparison across cell lines, expression values were z-scored per gene across all included cell lines. For each cell line, state-specific scores were computed by taking the mean z-scored expression across all genes assigned to each of the seven transcriptional states, yielding a seven-dimensional state score profile per cell line. To summarize differentiation along a single axis, we computed a melanocytic-like score as the mean of the melanocytic, transitory-melanocytic, and transitory state scores, and an undifferentiated/neural crest-like score as the mean of the neural crest-like, undifferentiated-neural crest-like, and undifferentiated state scores. The neural crest-like-transitory state was excluded from both aggregates. A one-dimensional differentiation axis was then defined as DiffAxis = (melanocytic program score) − (undifferentiated program score), where higher values indicate a more melanocytic-like transcriptional profile and lower values indicate a more undifferentiated/neural crest-like profile. To generate binary differentiation-state classes, DiffAxis values across all melanoma cell lines were fit with a two-component Gaussian mixture model using the mclust package[65]. Each cell line was assigned to a component based on posterior membership, and the component with the higher mean DiffAxis was labeled melanocytic-like, while the other was labeled undifferentiated/neural crest-like. An analytic decision boundary corresponding to the intersection of the component densities (accounting for mixture proportions) was used as the classification threshold.

### Partial least squares discriminant analysis (PLS-DA)

We used partial least-squares discriminant analysis (PLS-DA) to test whether melanocytic-like and undifferentiated/neural crest-like melanoma cell lines could be distinguished based on their epigenetic dependency profiles. Analysis was performed on the 51 CCLE melanoma cell lines with matched DepMap gene dependency data (34 melanocytic-like; 17 undifferentiated/neural crest-like) using dependency scores for 735 curated epigenetic regulators as input features. To account for class imbalance, conditional down-sampling was applied in iterations to limit the maximum class ratio to 3:1, using fixed random seeds for reproducibility. The optimal number of latent variables was selected by maximizing the cross-validated area under the receiver operating characteristic curve (ROC AUC) using 5-fold cross-validation. Model performance was summarized across 100 independent down-sampling iterations and reported as mean ROC AUC ± SD. Variable importance in projection (VIP) scores were computed using the first two latent variables and assigned directionality based on logistic regression coefficients fit on the same preprocessed input matrix. Signed VIP scores were used to rank epigenetic dependencies most associated with each differentiation state, with |VIP| > 1 used as the threshold for state-associated features.

### Pan-cancer complex dependency analysis from DepMap gene dependency profiles

Genome-wide DepMap CRISPR gene dependency probability scores for CCLE cancer cell lines were downloaded from the DepMap Portal (DepMap Public 25Q2 release; file CRISPRGeneDependency.csv) and analyzed in R. Gene dependency scores were represented on a probability-of-dependency scale ranging from 0 to 1, where 0 indicates no dependency and 1 indicates complete dependency (i.e., loss of viability upon gene perturbation), based on Chronos-processed CRISPR gene effect estimates[27]. Cell lines were annotated with cancer lineage using CCLE Model metadata (Oncotree primary disease/lineage). Non-cancerous lineages were excluded prior to downstream modeling. To ensure adequate representation per cancer type, lineages with fewer than five cell lines were removed. In total, 76 lineages were present prior to filtering, and 42 lineages were retained. Analyses were restricted to a filtered set of 735 epigenetic regulator genes from EpiFactors[28], representing genes found in the DepMap dependency matrix and containing no missing (NA) values across cell lines.

To identify lineage-enriched dependencies at the single-gene level, one-versus-rest differential dependency analysis was performed separately for each lineage using limma[29]. A design matrix was constructed with one indicator variable per lineage (no intercept), and for each lineage, a contrast was defined comparing that lineage against the mean of all other retained lineages. Linear models were fit per gene across cell lines and empirical Bayes moderation was applied to shrink variance estimates. For each gene *g* and lineage *ℓ*, the effect size (log-fold change; logFC) was defined as the difference between the mean dependency in lineage ℓ and the mean across all other lineages:

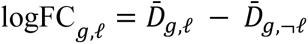

where *D̄_g,ℓ_* denotes the mean dependency score for gene *g* across cell lines in lineage ℓ, and *D̄_g,¬ℓ_* denotes the mean across all other retained lineages. Significance was assessed using Benjamini-Hochberg false discovery rate (FDR) correction across genes per contrast. Genes were considered lineage-enriched when FDR ≤ 0.05 and logFC ≥ 0.10, where positive logFC indicates greater dependency in the lineage compared to the aggregate of other lineages.

Chromatin regulatory complexes were initially compiled from EpiFactors. To generate a compact and interpretable complex list suitable for downstream analysis, we performed a manual curation and collapsing procedure to remove redundancy and harmonize naming across closely related complexes. First, complex names were standardized by resolving synonyms and alternative naming conventions to a single canonical label. Second, sub-complexes, assembly-specific variants, and highly overlapping entries describing the same biological machinery were merged into unified parent categories (e.g., SWI/SNF variants were consolidated into canonical BAF/cBAF, PBAF, and non-canonical ncBAF). Third, duplicate entries corresponding to the same complex listed under multiple subunit targets were collapsed into a single representative complex. This procedure produced a reduced, non-redundant complex reference set containing 43 complexes spanning 216 unique genes. Complex sizes ranged from 2 to 19 genes per complex (median = 6; mean = 6.67).

To assess lineage-enriched complex dependencies, gene dependency scores were aggregated into cell line-level complex dependency scores and tested using the same limma one-versus-rest framework. For each cell line *i* and complex *c*, complex dependency score was computed by averaging dependency scores across all complex member genes:

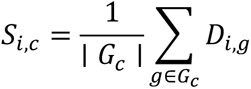

where *D_i_*_,g_ is the dependency score for gene *g* in cell line *i*, and *G_c_* is the set of genes annotated to complex *c*. The resulting complex score matrix {*S_i,c_*} was then analyzed using limma with the same design matrix and one-versus-rest contrasts described above (lineage as the grouping variable), fitting linear models per complex followed by empirical Bayes moderation. For each complex *c* and lineage *ℓ*, the effect size was defined analogously as

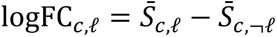

where *S̄_c,ℓ_* is the mean complex dependency score within lineage *ℓ* and *S̄_c,¬ℓ_* is the mean across all other retained lineages. Complexes were considered lineage-enriched when FDR ≤ 0.10 and logFC > 0, where positive logFC indicates greater dependency in the lineage compared to the aggregate of other lineages.

The nominal FDR ≤ 0.10 threshold applied in the complex-level analysis was intentionally more permissive than the stringent FDR ≤ 0.05 threshold used in the gene-level analysis, because complex-level dependency scores were calculated by averaging the dependency scores of all annotated subunits within a complex, including subunits that may not individually contribute to a lineage-enriched dependency. This averaging can dilute true effect sizes and reduce statistical power, justifying the use of a more permissive threshold at this stage as an exploratory screen for candidate epigenetic complexes, which subsequently corroborated by independent gene-level analyses at FDR ≤ 0.05, followed by gene set enrichment analysis.

Hierarchical clustering and heatmap visualization were performed using lineage-by-feature matrices of limma logFC values (features representing either genes or complexes). Matrices were transformed by column-wise z-scoring to emphasize relative enrichment patterns across features for each complex. Hierarchical clustering was performed for both rows and columns using Euclidean distance and the complete method.

### Statistical analysis and reproducibility

Sample sizes for cell line-based analyses were determined by the availability of publicly accessible datasets (DepMap and CCLE). For experimental approaches, no statistical method was used to predetermine sample size. For single-cell covariate analyses, the number of cells visualized per condition was selected based on similar quantitative imaging studies in the literature. Comparisons of dependency scores between groups (e.g., one lineage versus all other lineages or CXXC1^High^ versus CXXC1^Low^ melanoma subgroups) were performed using two-sided Wilcoxon rank-sum tests, as these data did not assume normal distributions.

For single-cell immunofluorescence experiments, differences in log-transformed fluorescence intensity between si-CXXC1 and si-NTC conditions were assessed using two-sided Welch’s t-tests on replicate-level summary statistics (mean log-transformed fluorescence intensity per biological replicate; n = 3 replicates per condition). For percentage-based analyses (% CXXC1^High^ cells, % H3K4me3^High^ cells, normalized cell count, and % Ki-67^High^ cells), comparisons between si-CXXC1 and si-NTC conditions were performed using linear mixed-effects models, with treatment as a fixed effect and well as a random effect (n = 3 biological replicates or wells per condition). One-sided P values were adjusted for multiple comparisons across cell lines for each marker using the Benjamini-Hochberg method. For these measurements, interaction analyses were additionally performed by extending the model to include cell line (or cell-line group) and its interaction with treatment. Satterthwaite F-tests were used to determine whether the effect of CXXC1 knockdown differed significantly across cell lines or cell-line groups. Effect sizes were expressed based on the percent reduction relative to si-NTC, calculated as 100 × (1 − e^β^), where β represents the model-estimated log-scale difference between si-CXXC1 and si-NTC.

Bar graphs display mean ± SEM across biological replicates. Boxplots show the median as the center line and the interquartile range (IQR) as the box, with whiskers extending to 1.5×IQR. In violin-boxplots, the violin represents the full distribution of the measurement (kernel density estimate), with an embedded boxplot indicating the median and IQR. Linear regression analyses were performed using ordinary least-squares regression, and coefficient of determination (R^2^) was calculated to quantify goodness of fit. Unless otherwise specified, all statistical tests were two-sided, and adjusted P values (FDR) were reported where multiple comparisons were performed. All statistical analyses were conducted in R (version 4.x).

## Data and code availability

RNA sequencing data collected in this study have been deposited in Gene Expression Omnibus (https://www.ncbi.nlm.nih.gov/geo/) with the accession number GSE324132. The data and original code for data analysis performed in this paper are publicly available at GitHub: https://github.com/fallahi-sichani-lab/chromatin-complex-dependencies-Set1C-COMPASS-melanoma. This paper analyzes existing, publicly available data, and the information for the public datasets are described in Methods. Any additional information required to reanalyze the data reported in this paper is available upon request.

## Author Contributions

**Conceptualization:** Luisa Quesada Camacho, Mohammad Fallahi-Sichani

**Data Curation:** Luisa Quesada Camacho

**Formal Analysis:** Luisa Quesada Camacho

**Funding Acquisition:** Mohammad Fallahi-Sichani

**Investigation:** Luisa Quesada Camacho

**Methodology:** Luisa Quesada Camacho, Mohammad Fallahi-Sichani

**Software:** Luisa Quesada Camacho

**Supervision:** Mohammad Fallahi-Sichani

**Visualization:** Luisa Quesada Camacho, Mohammad Fallahi-Sichani

**Writing:** Luisa Quesada Camacho, Mohammad Fallahi-Sichani

## State of Competing Interests

The authors declare that they have no competing interests.

## Acknowledgments

We thank members of the Fallahi-Sichani laboratory for their technical contributions, helpful suggestions, and discussion. We acknowledge the University of Virginia Genome Analysis and Technology Core (RRID: SCR_018883) and Research Computing at the University of Virginia for providing technical support and computational resources. This work was supported by NIH grants R01-CA249229 and P30-CA044579 (University of Virginia Cancer Center Support Grant).

## Supplementary Information

**Figure S1. Set1C/COMPASS dependency in melanoma cell lines is not explained by genetic alterations in, or expression of, Set1C/COMPASS subunits, or abundance of related global histone post-translational modifications. (A, B)** Pearson’s correlation analysis of CXXC1 dependency with SETD1A (A) and SETD1B (B) dependency scores across 67 CCLE melanoma cell lines. **(C)** Mutation status of genes encoding Set1/COMPASS subunits across melanoma cell lines (non-silent mutation = Mutant; silent/no mutation = WT). Cell lines are ordered by DepMap CXXC1 dependency score. **(D)** Comparison of CXXC1 dependency scores between melanoma cell lines classified as Set1/COMPASS mutant versus wild-type (WT), where “mutant” indicates at least one non-silent mutation in any Set1C/COMPASS subunit. P values were calculated using a two-sided Wilcoxon rank-sum test. **(E)** Comparison of Set1C/COMPASS subunit expression levels between subgroups of melanoma cell lines stratified by CXXC1 dependency (CXXC1^Low^ vs CXXC1^High^; threshold = 0.4). Boxplots indicate the median and interquartile range, and the Wilcoxon rank-sum P value and group sample sizes (n) are shown. **(F)** Correlation between CXXC1 dependency score and global histone post-translational modification (PTM) levels across melanoma cell lines with available CCLE Global Chromatin Profiling data (n = 34). For each of 42 histone PTMs, the Pearson’s correlation coefficient (r) between CXXC1 dependency and PTM abundance was plotted against -log_10_ (P value). PTMs reaching the significance threshold (P ≤ 0.05) are highlighted in red and labeled; all other PTMs are shown in gray.

**Figure S2. Melanoma-enriched Set1C/COMPASS dependency is not restricted to a specific differentiation state. (A)** Two-component Gaussian mixture modeling of the melanoma differentiation axis used to assign melanocytic-like versus undifferentiated/neural crest-like classes. Histogram shows the distribution of differentiation axis values across 71 melanoma cell lines; overlaid curves show the fitted component densities. The red dashed line denotes the decision boundary at the intersection of the two component densities, used as the classification threshold. **(B)** ROC curves summarizing PLS-DA performance for classifying melanoma cell lines as melanocytic-like versus undifferentiated/neural crest-like based on epigenetic gene dependency profiles. Model performance was evaluated by five-fold stratified cross-validation; curves show ROC performance across cross-validation runs, and the mean ± SD area under the curve (AUC) is reported.

**Figure S3. CXXC1 depletion reduces H3K4me3 in CXXC1-dependent melanoma cell lines. (A-F)** Quantitative changes in CXXC1, H3K4me3 and Ki-67 protein levels induced by CXXC1 siRNA (si-CXXC1) relative to non-targeting control (si-NTC) across five melanoma cell lines. Log-transformed distributions of single-cell fluorescence intensities for each protein marker are shown with boxplots indicating the median signal and interquartile range for each condition (A, C, E). Horizontal dashed lines mark high/low thresholds for single-cell intensities derived from two-component Gaussian mixture models fit to each signal (B, D, F). P values for single-cell distribution plots compare conditions using two-sided Welch’s t-tests on per-replicate mean log-intensities (n = 3 replicates per condition). **(G, H)** Quantitative changes in markers of cell cycle progression, p-Rb^Ser807/811^ (G) and p-Rb^Thr373^ (H), induced by CXXC1 siRNA (si-CXXC1) relative to non-targeting control (si-NTC). Log-transformed distribution of single-cell level fluorescence intensities are shown with boxplots indicating the median signal and interquartile range for each condition. P values compare conditions using two-sided Welch’s t-tests on per-replicate mean log-intensities (n = 3 replicates per condition).

**Figure S4. CXXC1 depletion suppresses MYC- and E2F-linked transcription programs in H3K4me3-responsive cell lines. (A)** PC1 versus PC2 and PC3 scores capturing cell line-specific variations in gene expression. **(B)** siRNA-induced changes in CXXC1 transcript level across four cell lines as quantified by RNA sequencing analysis. (**C-D**) Pre-ranked gene set enrichment analysis of H3K4me3-responsive up- or down-regulated genes using KEGG pathways (C) and GO Biological Process terms (D).

**Figure S5. Association of Set1C/COMPASS dependency with MYC- and E2F-linked transcription programs across cancer lineages. (A-C)** Lineage-specific pre-ranked gene set enrichment analysis (GSEA; fgsea) for the Hallmark gene sets E2F Targets (A), MYC Targets V1 (B), and MYC Targets V2 (C), performed across all cancer lineages represented by 5 or more cell lines with DepMap gene-dependency profiles. For each cell line, Set1C/COMPASS dependency was defined as the mean dependency score of CXXC1, SETD1B, ASH2L, DPY30, RBBP5, WDR5, and WDR82. Within each lineage, genes were ranked by the Spearman correlation between their dependency scores and Set1C/COMPASS dependency across cell lines. The normalized enrichment score (NES) and FDR from pre-ranked GSEA for each lineage are shown on the x- and y-axes, respectively. Lineage-specific enrichment of Set1C/COMPASS dependency relative to other cancer lineages, as determined in Figure 1A, is represented by the z-scored mean dependency (marker color) and FDR (marker size). The following lineages with significantly enriched Set1C/COMPASS dependency are highlighted in red: melanoma (MEL), B-cell acute lymphoblastic leukemia (B-ALL), and head and neck squamous cell carcinoma (HNSC).

**Table S1. List of epigenetic gene dependencies associated with melanoma differentiation state identified by PLS-DA (|VIP| > 1).** Variable importance in projection (VIP) scores were computed from the PLS-DA model using the selected number of latent variables. Directionality was assigned using logistic regression coefficients fit on the same preprocessed dependency matrix, yielding signed VIP scores. Genes with |VIP| > 1 were retained as differentiation state-associated features. Positive signed VIP scores indicate higher association with the melanocytic-like state, whereas negative signed VIP scores indicate higher association with the undifferentiated/neural crest-like state.

**Table S2. Summary of statistical analysis results associated with the CXXC1 knockdown experiment performed in five melanoma cell lines. (A-C)** Statistical analysis of changes in the percentage of CXXC1^High^ cells (A), percentage of H3K4me3^High^ cells (B), and normalized cell count (C) following CXXC1 siRNA (si-CXXC1) treatment relative to non-targeting control (si-NTC) across five cell lines (n = 3 wells/biological replicates). Adjusted P values (FDR) were derived from linear mixed-effects models with treatment as a fixed effect and well/replicate as a random effect, using one-sided tests adjusted for multiple comparisons across cell lines using the Benjamini-Hochberg method. **(D)** Interaction analyses performed by extending the linear mixed-effects models to include cell line or cell-line group (H3K4me3-responsive versus H3K4me3-nonresponsive as determined in B) and their interaction with treatment. Satterthwaite F-tests were used to determine whether the effect of CXXC1 knockdown differed significantly across cell lines or cell-line groups.

