## Supplementary figures and images for "Systematic multivariate analysis of chromatin complex dependencies reveals Set1C/COMPASS as a melanoma-enriched epigenetic vulnerability"

### Supplemental Figure S1

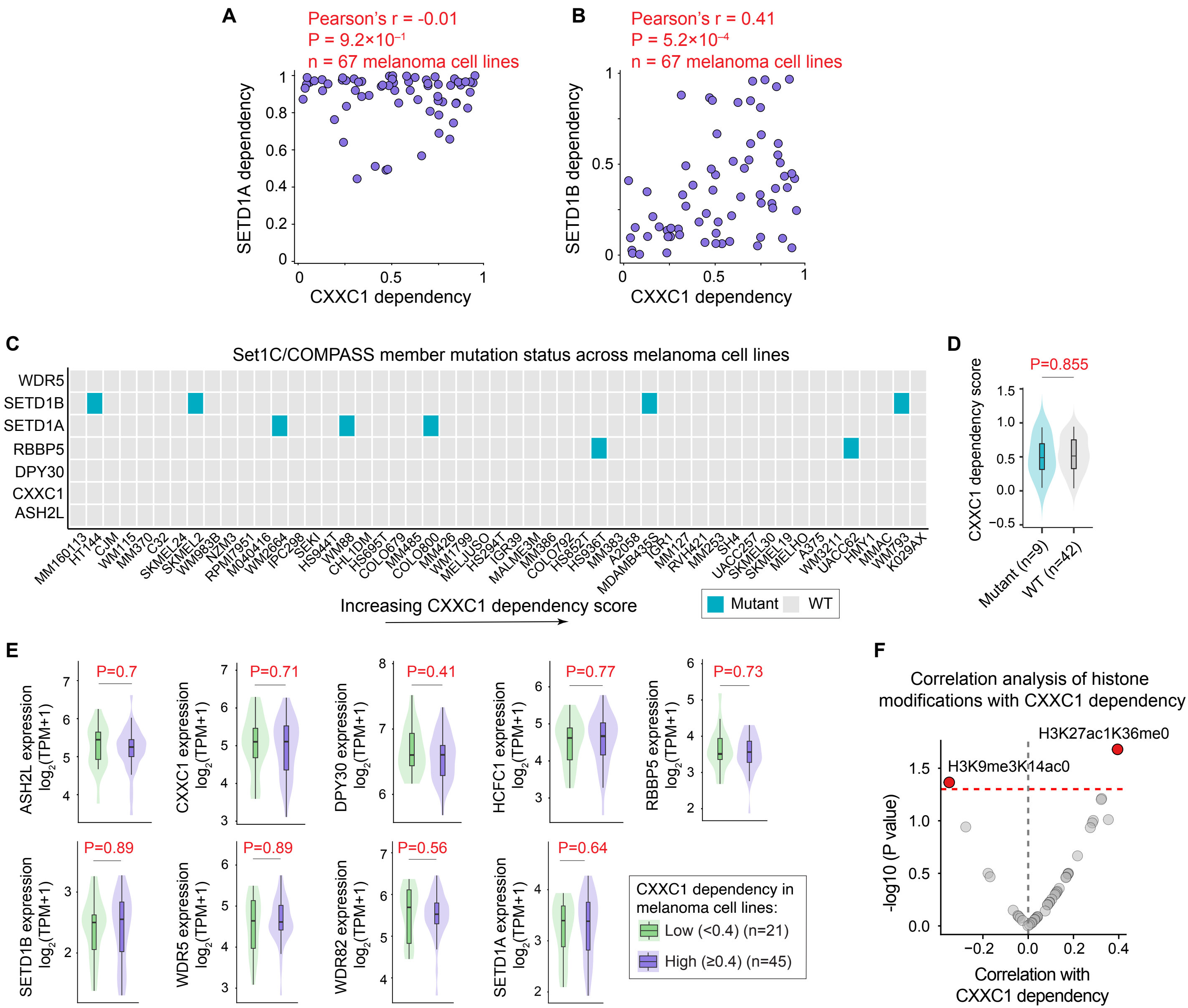

### Supplemental Figure S2

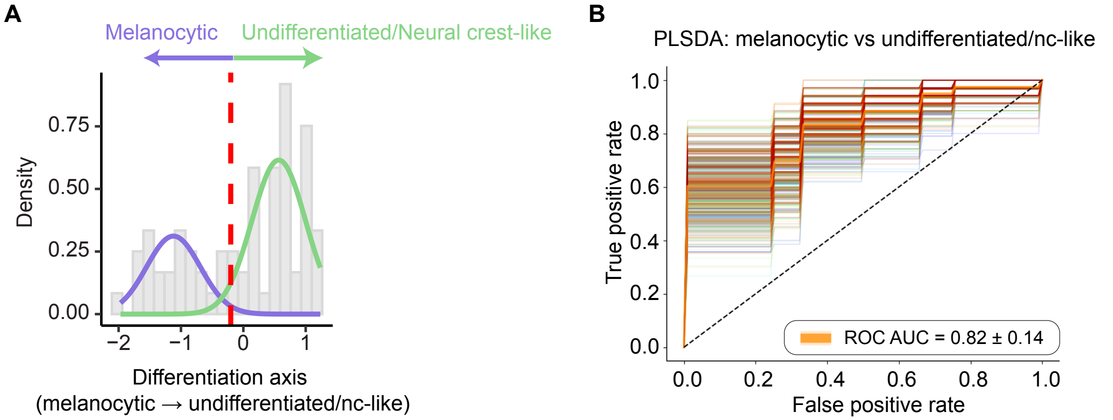

### Supplemental Figure S3

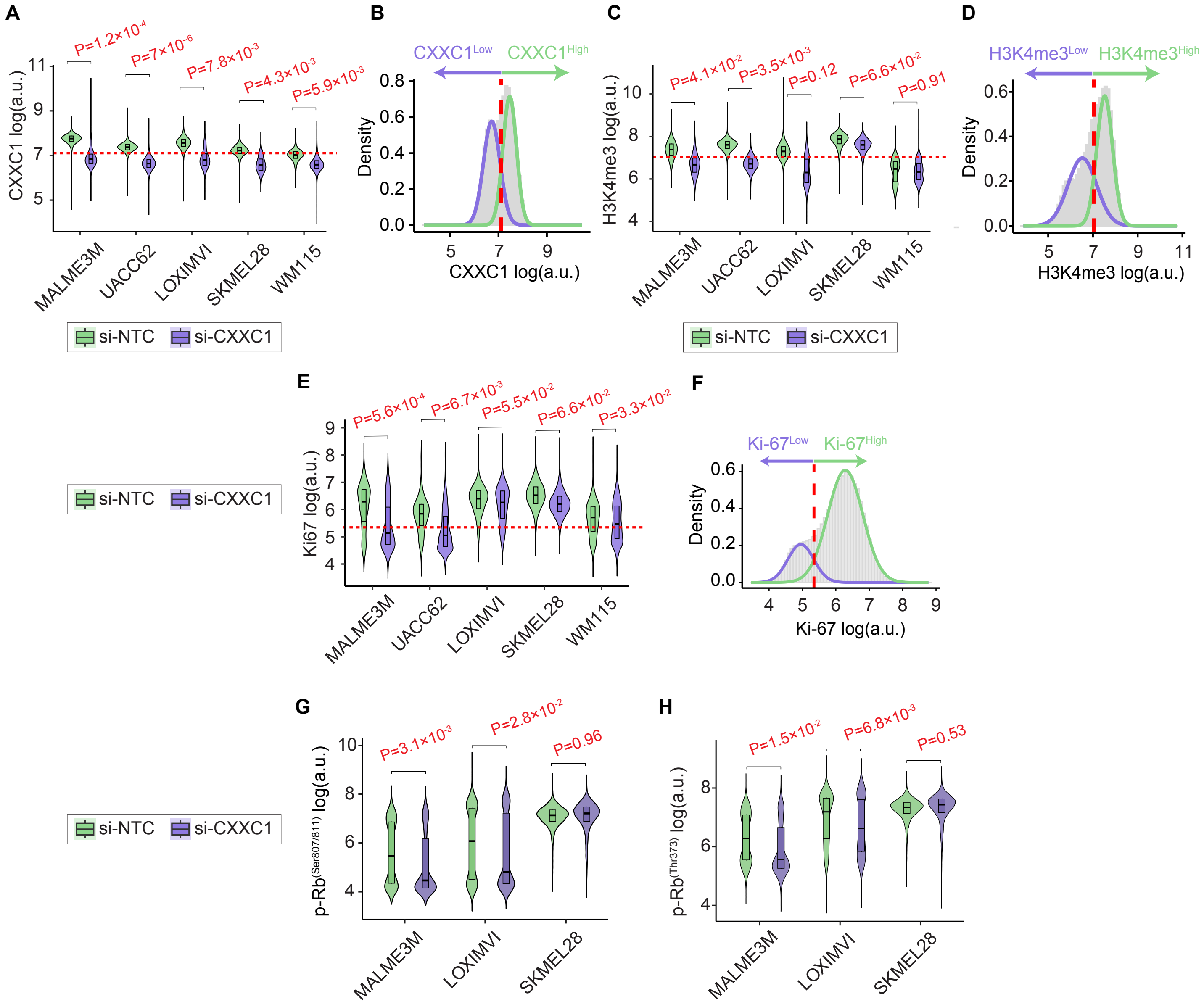

### Supplemental Figure S4

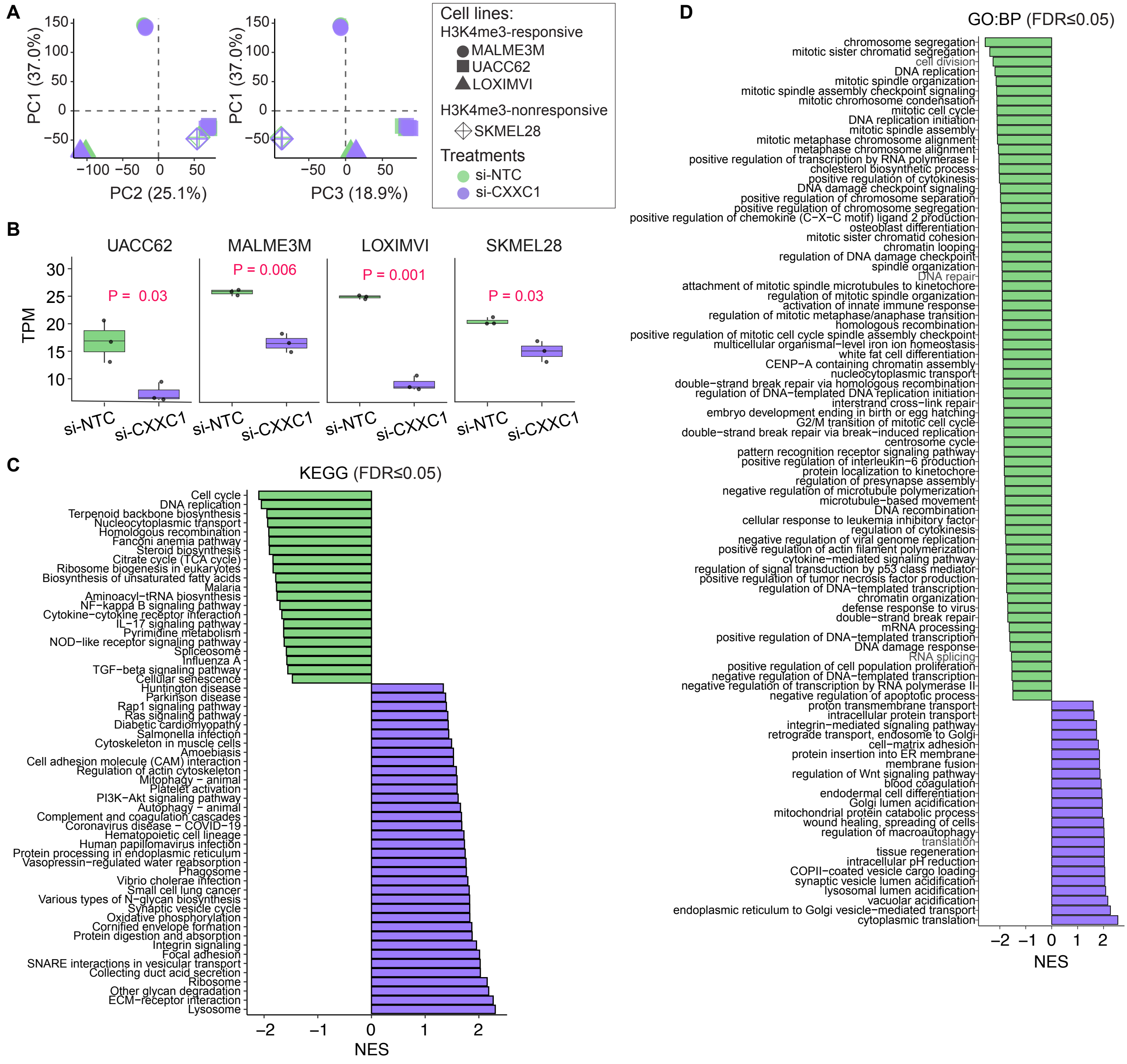

### Supplemental Figure S5

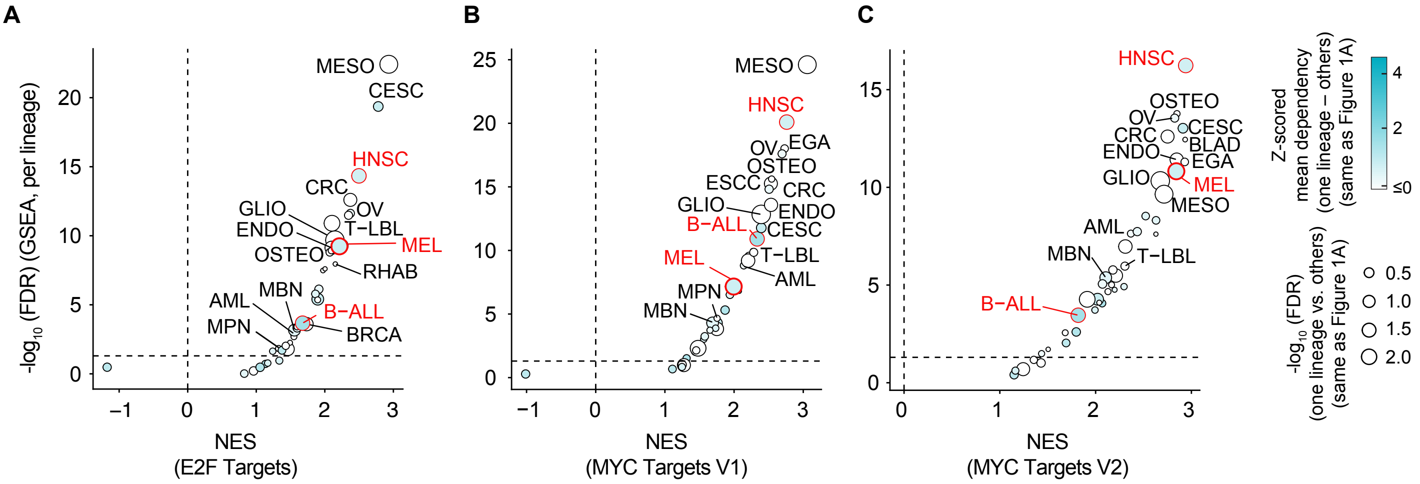
