## Supplemental Table S2 for "Systematic multivariate analysis of chromatin complex dependencies reveals Set1C/COMPASS as a melanoma-enriched epigenetic vulnerability"

**Table S2. Summary of statistical analysis results associated with the CXXC1 knockdown experiment performed in five melanoma cell lines.**

**(A)** Statistical analysis of changes in the percentage of $CXXC1^{High}$ cells following CXXC1 siRNA (si-CXXC1) treatment relative to non-targeting control (si-NTC) across five cell lines (n = 3 wells/biological replicates). Adjusted P values (FDR) were derived from linear mixed-effects models with treatment as a fixed effect and well/replicate as a random effect, using one-sided tests adjusted for multiple comparisons across cell lines using the Benjamini-Hochberg method.

| Measurement | Cell Line | % Reduction | Effect size (Cohen's d) | FDR | Outcome |
|---|---|---|---|---|---|
| % $CXXC1^{High}$ | MALME3M | 77.0% | 20.31 | 1.55e-3 | Significant CXXC1 depletion |
| % $CXXC1^{High}$ | UACC62 | 94.4% | 50.20 | 2.54e-6 | Significant CXXC1 depletion |
| % $CXXC1^{High}$ | LOXIMVI | 75.0% | 15.24 | 1.55e-3 | Significant CXXC1 depletion |
| % $CXXC1^{High}$ | SKMEL28 | 85.8% | 4.89 | 1.24e-2 | Significant CXXC1 depletion |
| % $CXXC1^{High}$ | WM115 | 86.2% | 5.99 | 5.04e-3 | Significant CXXC1 depletion |

**(B)** Statistical analysis of changes in the percentage of $H3K4me3^{High}$ cells following CXXC1 siRNA (si-CXXC1) treatment relative to non-targeting control (si-NTC) across five cell lines (n = 3 wells/biological replicates). Adjusted P values (FDR) were derived from linear mixed-effects models with treatment as a fixed effect and well/replicate as a random effect, using one-sided tests adjusted for multiple comparisons across cell lines using the Benjamini-Hochberg method.

| Measurement | Cell Line | % Reduction | Effect size (Cohen's d) | FDR | Outcome |
|---|---|---|---|---|---|
| % $H3K4me3^{High}$ | MALME3M | 73.2% | 3.50 | 1.63e-2 | H3K4me3-responsive |
| % $H3K4me3^{High}$ | UACC62 | 82.9% | 6.17 | 1.63e-2 | H3K4me3-responsive |
| % $H3K4me3^{High}$ | LOXIMVI | 74.1% | 2.20 | 4.75e-2 | H3K4me3-responsive |
| % $H3K4me3^{High}$ | SKMEL28 | 3.7% | 2.28 | 4.75e-2 | H3K4me3-nonresponsive |
| % $H3K4me3^{High}$ | WM115 | 10.0% | 0.13 | 4.42e-1 | H3K4me3-nonresponsive |

**(C)** Statistical analysis of changes in the normalized cell count following CXXC1 siRNA (si-CXXC1) treatment relative to non-targeting control (si-NTC) across five cell lines (n = 3 wells/biological replicates). Adjusted P values (FDR) were derived from linear mixed-effects models with treatment as a fixed effect and well/replicate as a random effect, using one-sided tests adjusted for multiple comparisons across cell lines using the Benjamini-Hochberg method.

| Measurement | Cell Line | % Reduction | Effect size (Cohen's d) | FDR | Outcome |
|---|---|---|---|---|---|
| Normalized cell count | MALME3M | 21.1% | 24.19 | 2.23e-5 | Significant reduction of cell count |
| Normalized cell count | UACC62 | 52.8% | 18.77 | 7.60e-5 | Significant reduction of cell count |
| Normalized cell count | LOXIMVI | 16.1% | 2.25 | 9.01e-2 | Modest reduction of cell count |
| Normalized cell count | SKMEL28 | -8.9% (increase by 8.9%) | -1.83 (increase by 1.83) | 9.39e-1 | No reduction in cell count |
| Normalized cell count | WM115 | 1.0% | 0.12 | 5.56e-1 | No reduction in cell count |

**(D)** Interaction analyses performed by extending the linear mixed-effects models to include cell line or cell-line group (H3K4me3-responsive versus H3K4me3-nonresponsive as determined in B) and their interaction with treatment. Satterthwaite F-tests were used to determine whether the effect of CXXC1 knockdown differed significantly across cell lines or cell-line groups.

| Measurement | Interaction | F | df1 | df2 | p-value |
|---|---|---|---|---|---|
| % $H3K4me3^{High}$ | siRNA condition × cell line | 5.35 | 4 | 20 | 4.27e-3 |
| % $H3K4me3^{High}$ | siRNA condition × cell line group | 5.26 | 1 | 26 | 3.01e-2 |
| Normalized cell count | siRNA condition × cell line | 34.85 | 4 | 20 | 9.42e-9 |
| Normalized cell count | siRNA condition × cell line group | 12.02 | 1 | 26 | 1.85e-3 |
